# Characterizing Physicochemical Selection in Protein Evolution with Property-Informed Models (PRIME)

**DOI:** 10.64898/2026.03.09.710461

**Authors:** Hannah Kim, Konrad Scheffler, Anton Nekrutenko, Darren P. Martin, Steven Weaver, Ben Murrell, Sergei L. Kosakovsky Pond

**Affiliations:** Institute for Genomics and Evolutionary Medicine, Temple University, Philadelphia, PA, USA; Department of Biology, Temple University, Philadelphia, PA, USA; Current address: Illumina, Inc., San Diego, CA, USA; Department of Biochemistry and Molecular Biology, The Pennsylvania State University, University Park, PA, USA; Institute of Infectious Disease and Molecular Medicine, University of Cape Town, Cape Town, South Africa; Department of Microbiology, Tumor and Cell Biology, Karolinska Institutet, Stockholm, Sweden

## Abstract

Standard probabilistic models of coding sequence evolution effectively identify where and when selection acts but remain agnostic to the mechanistic realization of these forces. We introduce PRIME (PRoperty Informed Models of Evolution), a framework of codon-level maximum likelihood methods—including global (G-PRIME), episodic (E-PRIME), and site-specific (S-PRIME) implementations—that explicitly model amino acid exchange-ability as a function of physicochemical properties. By parameterizing attributes such as molecular volume, hydropathy, and secondary structure propensities, PRIME resolves the biophysical basis of selective constraint across both the sequence and the phylogeny. At the site level, S-PRIME leverages an explicit biophysical taxonomy to precisely categorize residues as conserved, neutral, or changing for specific properties, resolving selective signals that remain invisible to traditional rate-based metrics. Our analysis of a benchmark of 24 diverse datasets and a genome-wide screen of 18,944 mammalian genes demonstrates that biophysical realism yields substantial improvements in model fit, acting synergistically with rate variation to explain complex evolutionary patterns. We find that power to detect physicochemical constraints at individual sites is fundamentally governed by simple informational redundancy (substitutions per unique amino acid; AUC = 0.91), with sensitivity exceeding 90% in data-rich alignments. E-PRIME reveals a distinct biophysical hierarchy: while core packing and beta-sheet scaffolds are rigidly conserved, alpha-helix propensity and surface electrostatics serve as the primary substrates for adaptive tuning. Furthermore, PRIME importance weights align with aspects of the primary semantic axes of deep learning representations (ESM-2) and capture key features of experimental fitness landscapes. By transforming abstract evolutionary rates into interpretable biophysical rules, PRIME provides a useful framework for characterizing the mechanistic drivers of protein diversity.

## 1 Introduction

Deciphering the selective pressures acting on viral genomes and other organisms requires robust probabilistic models of protein evolution (Durbin et al., 1998; Felsenstein, 2004; Yang, 2006). These statistical tools are indispensable for investigating phenomena such as host switching (Ngandu et al., 2009), immune response (Delport et al., 2008; Lacerda et al., 2010), and the emergence of drug resistance (Seoighe et al., 2007; Kosakovsky Pond et al., 2008; Murrell et al., 2012c). Yet, while current methodologies have achieved high sensitivity in detecting *where* in the sequence (Murrell et al., 2012a) and *when* in evolutionary history (Murrell et al., 2015) selection acts, they often fail to explain *how* this action is realized biochemically. A major gap remains in moving from the phenomenological detection of selection to the mechanistic characterization of the physicochemical rules that govern it.

The integration of physicochemical realism into codon substitution models traces its lineage to the pioneering work of Grantham (1974), who introduced a composite metric of amino acid differences based on composition, polarity, and volume. This concept was subsequently adapted by Miyata et al. (1979) and formalized by Goldman and Yang (1994), who scaled substitution rates using these fixed distances. Xia and Li (1998) further demonstrated that the genetic code itself minimizes polarity and hydropathy changes. Subsequent approaches attempted to categorize substitutions as property-altering or conserving (Sainudiin et al., 2005; Wong et al., 2006), a concept recently refined by the CoRa model’s distinction between conservative and radical changes (Weber and Whelan, 2019). Jones et al. (2020) introduced the phenotype-genotype branch-site model (PG-BSM), which explicitly links adaptive codon evolution to discrete phenotypic changes by detecting transient elevations in nonsynonymous rates, notably without requiring the non-synonymous/synonymous rate ratio *ω >* 1. Others, such as the LCAP model (Conant et al., 2007), sought to estimate property weights directly from data, though often at the cost of computational tractability or site-specific resolution. Despite these methodological advances, none have achieved widespread adoption; the vast majority of evolutionary analyses continue to rely on standard *ω* models agnostic of the amino-acids being exchanged, possibly because these alternatives lack the intuitive appeal of a simple rate ratio, have not been supported by robust, scalable software implementations, and have not been shown to significantly impact biologically meaningful inference.

Simultaneously, biophysical studies have underscored that evolutionary trajectories are shaped by fundamental physical constraints. Seminal work by Bloom, Wilke, and colleagues demonstrated that protein evolution is often governed by a global requirement to maintain thermodynamic stability, leading to a marginal stability regime where most mutations are destabilizing but neutral as long as the protein remains folded (Bloom et al., 2006, 2007). Mirny and Shakhnovich (1999) similarly showed that conserved positions often serve as nuclei for folding. These biophysical constraints—including folding kinetics and solubility—act as a universal filter on sequence space (Liberles et al., 2012; Serohijos and Shakhnovich, 2014). Furthermore, stability can promote evolvability by buffering the destabilizing effects of functionally innovative mutations (Tokuriki and Tawfik, 2009). However, the implicit assumption that protein evolution is primarily constrained by a rigid 3D fold constitutes a significant limitation for many existing approaches. A substantial fraction of the proteome consists of Intrinsically Disordered Regions (IDRs) and flexible linkers which lack stable tertiary structure yet play vital roles in signaling and regulation (Dunker et al., 2001). Evolution in these regions is distinct, often driven by the conservation of distributed features—such as net charge, hydropathy, or flexibility—rather than strict site-specific identities (Zarin et al., 2019). Standard substitution models, and even those grounded in structural thermodynamics, may fail to capture the relaxed or orthogonal constraints acting on these flexible segments.

In the past several years, the field has been revolutionized by deep learning approaches, including AlphaFold (Jumper et al., 2021) and protein language models (PLMs) like ESM (Rives et al., 2021; Lin et al., 2023). These models have achieved high success in learning the implicit grammar of protein sequences, effectively capturing high-order dependencies and structural constraints. However, they fundamentally trade mechanistic interpretability for predictive power. While a PLM might accurately predict *that* a specific substitution is unlikely, it operates as a high-dimensional black box, obscuring *why* that constraint exists. Whether a site is conserved due to electrostatic requirements, packing density, or an unknown functional dependency remains opaque. Consequently, the lack of an explicit biophysical grammar limits our ability to predict the impact of novel mutations, particularly in contexts like immune escape where viruses explore regions of sequence space not well-represented in training data.

To bridge this divide, we introduce PRIME (PRoperty Informed Models of Evolution), a comprehensive family of probabilistic models that filters evolutionary action through the prism of physicochemical properties. We present three complementary implementations: a G-PRIME (Global PRIME) model for characterizing gene-wide constraints, an E-PRIME (Episodic PRIME) framework for detecting episodic property selection, and an S-PRIME (Site-level PRIME) inference method for resolving residue-specific biophysics. By encoding the intuition that properties like polarity and volume are conserved or driven to change, we estimate exchangeabilities within a maximum likelihood framework. This approach yields a marked improvement in model fit and provides a formal, standalone platform for testing biophysical hypotheses. While PRIME can effectively decode the latent constraints of deep learning models, its primary power lies in its ability to explicitly resolve the biochemical mechanisms of evolution—transforming abstract substitution rates into concrete physical rules.

## 2 Methods

### 2.1 Model Description

We use the standard continuous-time Markov description of coding-sequence evolution (Goldman and Yang, 1994; Muse and Gaut, 1994). The transition matrix for finite time *t* ≥ 0 is given by *e^Qt^*; the rate matrix *Q* = {*q_ij_*} describes the instantaneous rate of substitution of codon *i* with codon *j*:

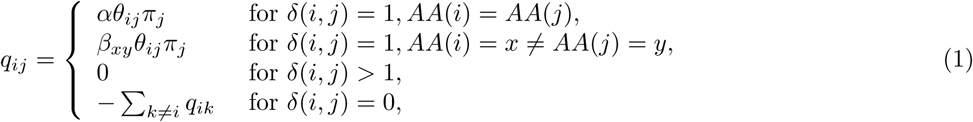

where *δ*(*i, j*) counts the number of nucleotide differences between codons *i* and *j*, and *AA*(*i*) stands for the amino acid corresponding to codon *i*. *θ_ij_* are the nucleotide-level mutation parameters implementing the general time reversible (GTR) model for nucleotide substitution biases, with the rate of transitions between Adenine and Guanine set to 1 to ensure model identifiability. *π_j_* are equilibrium codon frequencies. *α* is a multiplier for synonymous substitutions. *β_xy_* is a multiplier for nonsynonymous substitutions representing the exchangeability of amino acids *x* and *y*.

We depart from the standard model in that the nonsynonymous substitution rate depends on the amino acids involved in the substitution. Given a set of *D* properties such that the value of property *i* for amino acid *x* is *x_i_*, we model *β_xy_* as:

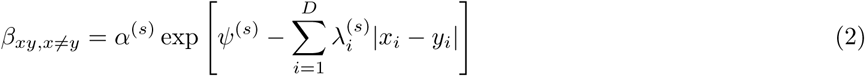

Here, *ψ*^(*s*)^ represents the baseline log nonsynonymous/synonymous rate ratio at site *s*. This formulation ensures that if all property weights 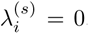, the model reduces to the standard MG94 model with *ω_s_*= exp(*ψ*^(*s*)^). The parameters 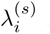 represent the importance of property *i* at site *s*. Positive values indicate conservation (purifying selection), where exchangeability declines as property difference increases. Negative values 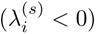 imply positive selection, where it is important for a property to change.

This general framework is deployed across three distinct implementations to capture selective constraints at different scales:

1. G-PRIME: Assumes constant *λ* values across the entire alignment, estimating the average physicochemical constraints acting on the gene product.
2. E-PRIME: Models *λ* as a random effect varying across phylogenetic branches and sites, allowing for the detection of episodic selection on specific properties.
3. S-PRIME: Estimates independent *λ* vectors for each codon site, resolving the fine-grained biophysical architecture of the protein. This functional form admits a direct interpretation in population genetics. The substitution rate is proportional to the probability of fixation of a new mutation, which, under strong purifying selection, scales exponentially with the selection coefficient *s* (specifically *P_fix_* ∝ *e*^2*N*^*^es^*). We posit that the effective selection coefficient for a substitution from amino acid *x* to *y* is composed of a baseline effect *s*_0_ (captured by *ψ*^(*s*)^) and specific physicochemical costs: 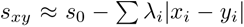. In this framework, 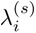 serves as a proxy for the product of the effective population size *N_e_* and the selective constraint intensity per unit of property change.

### 2.2 Amino Acid Properties

Models of this form can be used with different sets of amino acid properties. We consider primary properties such as chemical composition, polarity, volume, isoelectric point, and hydropathy (Conant et al., 2007), as well as composite property sets derived from dimensionality reduction techniques (Atchley et al., 2005). Within HyPhy, all property vectors are internally normalized to have zero mean and unit variance to ensure that the estimated *λ* coefficients are comparable across properties and to mitigate the impact of disparate scales on collinearity. The choice of properties represents a trade-off between interpretability and comprehensive coverage. Primary properties, such as the Kyte-Doolittle hydrophobicity index (Kyte and Doolittle, 1982) or the Zamyatnin volume scale (Zamyatnin, 1972), are physically intuitive and allow selective pressures to be mapped directly to specific biochemical mechanisms. In contrast, propensities for alpha-helix or beta-sheet formation (Chou and Fasman, 1974) capture marginal backbone geometric constraints. While these continuous scales do not fully account for the context-dependent nature of secondary structure formation (e.g., terminal capping or proline-induced kinks), they provide a statistically tractable proxy for the localized structural preferences that filter sequence variation.

### 2.3 Hierarchical Property Models

We define a nested hierarchy of models, designated as 2-prop through 5-prop, which sequentially incorporate five fundamental physicochemical properties. This hierarchical approach allows us to systematically test the explanatory power of each additional property.

1. 2-prop (Hydrophobicity + Volume): This baseline model includes the Kyte-Doolittle hydrophobicity index (Kyte and Doolittle, 1982) and residue volume (Zamyatnin scale) (Zamyatnin, 1972). These two properties represent the most fundamental constraints on protein folding: the hydrophobic effect drives the formation of the protein core, while steric constraints (volume) dictate packing density and surface accessibility. They are broadly complementary, as a residue can be small and hydrophobic (e.g., Valine) or large and hydrophobic (e.g., Tryptophan).
2. 3-prop (Adding Isoelectric Point): The 3-prop model adds the isoelectric point (pI), introducing charge as a distinct constraint. Electrostatic interactions are critical for protein stability, solubility, and specific ligand binding (Conant et al., 2007). While hydrophobicity governs the core, charge often dominates surface interactions and active sites, offering a dimension of constraint orthogonal to packing and burial.
3. 4-prop (Adding Alpha-Helix Propensity): We extend the model by including alpha-helix propensity (Chou and Fasman, 1974). This property captures local backbone geometric constraints essential for secondary structure formation. Helical propensity is distinct from simple volume or hydrophobicity, as it depends on specific side-chain branching and hydrogen bonding capabilities.
4. 5-prop (Adding Beta-Sheet Propensity): The full 5-prop model incorporates beta-sheet propensity (Chou and Fasman, 1974), completing the description of major secondary structure determinants. This allows the model to differentiate between regions constrained by different folding motifs, providing a more comprehensive physicochemical profile of the evolutionary landscape.

These models enable a suite of systematic testing procedures. We can employ model selection criteria to determine the optimal level of complexity for a given gene or dataset; we use the small-sample corrected AICc (Sugiura, 1978; Hurvich and Tsai, 1989) throughout this manuscript. These criteria allow us to effectively ask: How much physicochemical detail is needed to explain the observed evolution? Furthermore, Likelihood Ratio Tests (LRTs) can be used to statistically assess the significance of adding specific properties (e.g., testing 3-prop vs. 2-prop to isolate the importance of charge), thereby building a detailed map of the specific biochemical forces shaping protein diversity.

### 2.4 Incorporating Branch-Site Heterogeneity

To capture the complexity of evolutionary processes, we employ a branch-site random effects framework ported from the BUSTED method (Murrell et al., 2015). This approach allows the importance of physicochemical properties (*λ* parameters) to vary not only across sites but also across different lineages (branches) of the phylogenetic tree. By modeling the distribution of *λ* as a random effect, we can detect episodic selection on properties —instances where specific physicochemical constraints are relaxed or intensified on particular branches. This is crucial for identifying transient evolutionary events, such as adaptation to a new host or immune escape, which may not be evident under models that assume constant preferences across the entire phylogeny. This framework provides a flexible and robust means to infer the dynamic landscape of physicochemical constraints in protein evolution.

### 2.5 PRIME Testing Procedure for Sites

To resolve the biophysical drivers of evolution at single-residue resolution, we extend the PRIME framework to estimate and test site-level importance weights 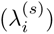. Analogous to the Fixed Effects Likelihood (FEL) method for *dN/dS* (Kosakovsky Pond and Frost, 2005), we assume that physicochemical constraints are temporally constant across the phylogeny at a given site but are modeled as completely independent parameters between sites. This site-by-site inference proceeds in two stages to balance computational efficiency with statistical precision. First, we estimate global nuisance parameters—including relative branch lengths, GTR nucleotide substitution rates, and equilibrium frequencies—jointly across the entire alignment. These parameters are subsequently fixed to reduce the dimensionality of the site-level optimization problem. In the second stage, we independently fit the site-specific parameters: the synonymous rate multiplier (*α*), the baseline log nonsynonymous/synonymous rate ratio (*ψ*^(*s*)^)—as opposed to *ω*^(*s*)^—and the vector of property importance coefficients 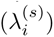 for each site *s*.

Statistical significance is assessed through a nested likelihood ratio test (LRT) framework. At each site *s*, we perform a set of *M* + 1 hypothesis tests, where *M* is the number of modeled properties (e.g., *M* = 5 for the 5-prop model):

1. Global Property Test: We first evaluate the omnibus null hypothesis that evolution at site *s* is independent of all modeled physicochemical properties 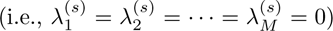. This is an LRT with *M* degrees of freedom.
2. Individual Property Tests: If the omnibus test rejects the null, we proceed to dissect the specific drivers by testing each property *i* individually. The null hypothesis 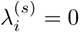 is assessed using a 1-degree-of-freedom LRT, determining whether property *i* contributes a statistically significant improvement to the model fit.

Given the large number of tests (*N* sites ×(*M* + 1) properties), and the expectations that many or most sites could have null results, stringent multiple testing correction is essential. We employ the hierarchical procedure described by Benjamini and Bogomolov (2014) to control the selective False Discovery Rate (sFDR). This two-stage approach first applies the Benjamini-Hochberg procedure to the *N* Global Property Tests (e.g., at FDR ≤ 0.05) to identify sites with significant biophysical structure. Only within this subset of *R* significant sites do we evaluate the Individual Property Tests. To account for selection bias (the cherry-picking of interesting sites), we apply the Holm-Bonferroni method at an adjusted significance level of *α* · *R/N* . This ensures strong control of the Family-Wise Error Rate (FWER) within each selected site while maintaining global FDR control, effectively mitigating the risk of false mechanistic attributions.

### 2.6 Structural Analysis

To map evolutionary constraints to physical structure, we integrated site-level statistics with three-dimensional protein coordinates for 21 of the 24 benchmark datasets (Table S2). For each dataset, we generated a consensus amino acid sequence from the translated multiple sequence alignment and aligned it to the corresponding PDB chain using the Smith-Waterman local alignment algorithm (Biopython PairwiseAligner, parameters: mode=’local’, open gap score=−10, extend gap score=−0.5). This protocol ensured robust site-to-structure mapping, even when the experimental structure represented a partial fragment or a distant homolog of the analyzed sequence. We extracted key structural features for each mapped site using the Biopython PDB module. Solvent accessibility was computed via the Shrake-Rupley algorithm (Shrake and Rupley, 1973) and normalized to Relative Solvent Accessibility (RSA) using theoretical maximum SASA values (Tien et al., 2013). Secondary structure assignments were generated using the DSSP algorithm (Kabsch and Sander, 1983). To capture the local packing environment, we also calculated normalized B-factors (chain-specific Z-scores), the contact number within a 10° A radius (*CN*_10_, defined by C*α*–C*α* distances), and polypeptide backbone torsion angles (*φ* and *ψ*). These features provide the structural context necessary to interpret the spatial distribution of physicochemical constraints.

While we utilize static PDB structures as a representative snapshot of each protein’s native fold, we acknowledge that residues exist within dynamic conformational ensembles. A single RSA value may not capture the full range of solvent exposure or packing density experienced by a residue during its functional cycle. Furthermore, structural features were mapped to monomeric PDB chains. While it is true that biological assemblies (e.g., the HA trimer) mask additional surface area at interfaces, the use of monomeric RSA provides a consistent measure of ”intrinsic” buriedness across our taxonomically diverse benchmark. The use of static reference structures remains the standard for large-scale structural-evolutionary mapping.

### 2.7 Comparison with protein language models

To investigate the relationship between explicit physicochemical constraints and the implicit representations learned by deep learning, we benchmarked site-specific PRIME importance weights (*λ*) against residue embeddings from the ESM-2 protein language model (Lin et al., 2023). For each dataset, we computed per-residue embeddings using the esm2 t33 650M UR50D architecture (layer 33, 1280 dimensions) for the primary reference sequence. A critical methodological step involved the precise alignment of these residue-level embeddings (*L_seq_*) with the alignment-level PRIME inferences (*L_aln_*). We mapped each non-gap residue in the reference to its corresponding MSA column, filtering the results to include only those sites present in the reference structure. To distill the high-dimensional PLM representation into interpretable components, we performed Principal Component Analysis (PCA) on the aligned ESM embeddings for each dataset. We then quantified the alignment between the two frameworks by calculating the Spearman rank correlation (*ρ*) between site-specific PRIME property weights and the top principal components of the ESM-2 embeddings. Following Rives et al. (2021), who established that the primary axes of PLM latent spaces frequently track fundamental biophysical features, we defined the recoverability of a property as the maximum absolute correlation between that property’s weight and any of the top five principal components. This approach formally tests whether the black-box predictions of the PLM are capturing the same underlying biophysical signal as the explicit PRIME model, providing a quantitative bridge between statistical phylogenetics and representation learning.

### 2.8 Genome-wide Screening Protocol

To characterize the landscape of physicochemical constraints across the mammalian proteome, we analyzed a high-quality subset of coding sequences from the 120-mammal genome alignment (Hecker and Hiller, 2019). The initial collection was filtered to ensure robust phylogenetic inference, retaining only alignments that met the following criteria: (1) contained at least 10 sequences; (2) had a coding length of at least 50 codons; and (3) exhibited a total tree length between 0.5 and 50 substitutions per site. This procedure yielded a final dataset of 18,944 gene alignments. For each gene, we fitted a suite of evolutionary models: the standard MG94 baseline, the CoRa model (Weber and Whelan, 2019), and the G-PRIME and E-PRIME implementations of the 5-prop PRIME model. All model fits were performed using HyPhy v2.5.94 or later. We employed the 5-property parameterization (Hydrophobicity, Volume, Isoelectric Point, Alpha-Helix Propensity, Beta-Sheet Propensity) as the default for this screen, based on its superior performance in our preliminary benchmarks. Statistical significance of property conservation or diversification was assessed using Likelihood Ratio Tests (LRTs) corrected for multiple testing via the False Discovery Rate (FDR) procedure (FDR ≤ 0.05). Genes where model fitting failed to converge for any of the comparison models were excluded from the final comparative analysis to ensure a strictly matched dataset.

### 2.9 Simulation Strategy for S-PRIME Performance Assessment

To systematically evaluate the statistical performance of S-PRIME in detecting site-specific physicochemical constraints, we designed a simulation framework that models the realistic scenario of localized biophysical pressure against a background of neutral evolution. Our approach assesses both the control of False Positive Rate (FPR) and the power to detect true positive signals under varying regimes of constraint intensity. For each of the 24 benchmark datasets, we first estimated global model parameters under the neutral null hypothesis (MG94), obtaining gene-specific estimates for branch lengths, nucleotide substitution biases (*θ*), and codon equilibrium frequencies (*π*). These parameters served as the seed for generating replicate alignments under two distinct conditions. First, to establish a baseline for Type I error control, we simulated alignments entirely under the null MG94 model (scenario null mg94), where all property importance weights are strictly constrained to zero (*λ* = 0). Second, to characterize detection power, we implemented a chimeric simulation strategy. We defined a set of over 20 selective scenarios representing diverse biophysical regimes. These include Conserved and Diversifying scenarios where a single focal property is constrained (e.g., *λ*_Vol_ = 3.0) while all other parameters are set to zero to ensure a clean background. We also included Empirical scenarios derived from clusters of site-level constraints observed in biological data, and Easy/Hard multi-property combinations. For each replicate, we generated two parallel alignments: a Null alignment evolved under MG94 and an Alternative alignment evolved under the 5-property G-PRIME model parameterized with the scenario-specific *λ* vector. We then constructed a spliced dataset by retaining the first 80% of sites from the Null alignment and appending the final 20% of sites from the Alternative alignment. This protocol injects a specific biophysical signal into a defined genomic window, allowing us to strictly quantify the method’s sensitivity to detect localized constraints while maintaining the phylogenetic context of the original gene. Power was calculated using the proportion of Alternative sites correctly identified as significant by the S-PRIME omnibus test after False Discovery Rate (FDR) correction (*q* ≤ 0.10).

### 2.10 Comparison with Experimental Fitness Landscapes

To validate the biological relevance of inferred physicochemical constraints, we benchmarked S-PRIME against Deep Mutational Scanning (DMS) data for the H3N2 Hemagglutinin protein (Lee et al., 2018). We integrated the A/Perth/16/2009 (H3N2) sequence used in the DMS assay into a historical H3N2 alignment (Bush et al., 1999) and performed a masked imputation analysis as follows. For each site in the Perth sequence, we discarded the observed codon and inferred the conditional probability distribution of all 61 sense codons given the phylogenetic history and the fitted model parameters (PRIME). This procedure parallels the masked language modeling objective used in deep learning.

For comparison, we generated zero-shot predictions using the ESM-2 protein language model (esm2 t33 650M UR50D) (Lin et al., 2023). We masked each position in the translated Perth amino acid sequence and extracted the probability distribution over the 20 standard amino acids. We evaluated the alignment between model predictions and experimental fitness using three metrics calculated over variable sites: (1) Pearson correlation, capturing the linear relationship between predicted probabilities and experimental preferences; (2) Spearman rank correlation, assessing the monotonicity of the ranking; and (3) Jensen-Shannon Divergence (JSD), quantifying the information-theoretic distance between the predicted and experimental distributions. Additionally, we defined a strongly preferred set for each site (residues with probability *>* 0.10) and calculated the Jaccard index and total probability mass coverage to assess the model’s ability to identify the subset of viable amino acids.

## Code and Data Availability

The PRIME framework is implemented as a module within the standard library of HyPhy (v2.5.94+). Source code for the inference engine is available at https://github.com/veg/hyphy. Supporting scripts and data for the analyses presented in this manuscript are available at https://github.com/veg/PRIME.

## 3 Results

### 3.1 Benchmarking Model Performance

To comprehensively evaluate the performance of PRIME and its branch-site extension (E-PRIME), we employed a suite of 21 diverse datasets previously established for model benchmarking (Lucaci et al., 2023). These datasets, whose functional and structural characteristics are detailed in Supplementary Table S1, represent a comprehensive cross-section of taxonomic scales, evolutionary rates, and biological functions, encompassing sequences from viruses (e.g., HIV-1, Influenza A, SARS-CoV-2, Flavivirus), mammals (e.g., primate lysozyme, camelid VHH, vertebrate rhodopsin), invertebrates (Drosophila Adh), plants (rbcL), and bacteria (Streptococcus PTS). This collection spans diverse selective regimes—from the intense diversifying selection of immune escape and sexual conflict to the stringent functional conservation of metabolic enzymes. To broaden the biophysical scope, we added three specialized datasets: Cytochrome C Oxidase I (COXI) for membrane protein constraints, Type I Collagen (COL1A1) for fibrous proteins with rigid repetitive motifs (Mu et al., 2021), and Amelogenin (AMELX) (Randall et al., 2022) to assess performance on intrinsically disordered proteins. We conducted a head-to-head comparison of four models (summarized in Table 2): (1) *MG*94×*REV*, the standard codon substitution baseline; (2) G-PRIME, incorporating alignment-wide property weights; (3) E-PRIME, the branch-site random effects extension; and (4) CoRa (Weber and Whelan, 2019), which we re-implemented within the Muse-Gaut framework to ensure a direct and fair comparison of rate matrices.

**Table 1:** Key parameters and symbols used in PRIME models. Parameters such as *α*, *ψ*, and *λ* may vary across sites and/or branches depending on the specific model configuration (global vs episodic vs site).

| Symbol | Description |
| --- | --- |
| $\alpha$ | Synonymous substitution rate multiplier |
| $\beta_{xy}$ | Nonsynonymous substitution rate multiplier between amino acids $x$ and $y$ |
| $\psi$ | Baseline log non-synonymous/synonymous rate ratio |
| $D$ | Number of modeled physicochemical properties (e.g., $D = 5$ for 5-prop) |
| $\lambda_i$ | Importance coefficient for the $i$th physicochemical property |
| $x_i$ | Value of the $i$th property for amino acid $x$ (e.g., $x_2 =$ value of property 2, volume, for amino acid $x$) |

**Table 2:**
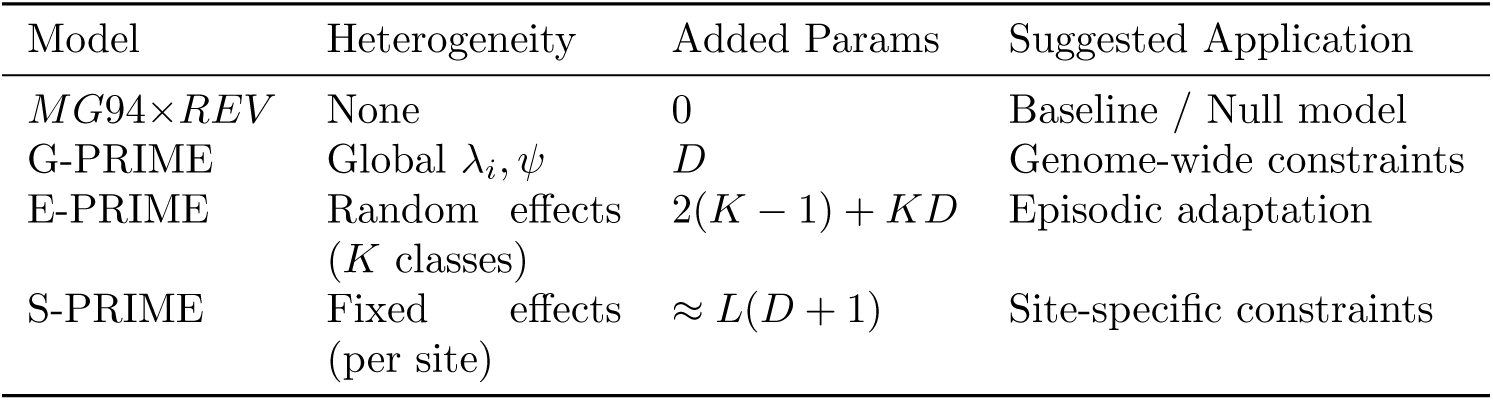
Architectural summary and parameterization of evaluated evolutionary models. PRIME configurations are distinguished by the level of selection heterogeneity they capture, ranging from genome-wide averages to site-specific biophysical drivers. We report the nature of heterogeneity, the number of additional parameters relative to the MG94+REV baseline (*D*: number of physicochemical properties; *K*: number of mixture components; *L*: number of alignment sites), and the intended biological applications for each model.

| Model | Heterogeneity | Added Params | Suggested Application |
| --- | --- | --- | --- |
| *MG94*×*REV* | None | 0 | Baseline / Null model |
| G-PRIME | Global $\lambda_i, \psi$ | $D$ | Genome-wide constraints |
| E-PRIME | Random effects<br>($K$ classes) | $2(K - 1) + KD$ | Episodic adaptation |
| S-PRIME | Fixed effects<br>(per site) | $\approx L(D + 1)$ | Site-specific constraints |

### 3.2 Substantial Improvements in Model Fit

As demonstrated in Supplementary Table S2, explicitly modeling physicochemical properties yields a substantial and consistent improvement in model fit across the benchmark. PRIME models outperformed both the standard *MG*94×*REV* baseline and the CoRa model in nearly every instance, proving that site-specific property constraints provide a far more nuanced and accurate description of evolution than fixed radical/conservative classifications. Notably, the magnitude of fit improvement is strongly coupled to the total evolutionary information content (*L* × *T*, *L* – alignment length, *T* – tree length, the product of which approximate the total expected number of substitutions); we observed a significant positive correlation between the expected number of substitutions and ΔAICc (Spearman *ρ* ≈ 0.65*, p <* 0.001). Large, divergent datasets such as rbcL and Mammalian Collagen exhibit substantial gains in fit (ΔAICc *>* 990). The 5-prop model emerged as the robust default choice for the majority of datasets, particularly when sufficient information is available to resolve complex constraints.

### 3.3 Dissecting Property-Specific Constraints

Beyond overall fit, S-PRIME enables a meticulous dissection of the specific biochemical properties that drive evolutionary trajectories. We summarized the global importance weights (*λ*) across the benchmark datasets in Figure 1 (raw values provided in Supplementary Table S3). Positive *λ* values (blue) denote properties subject to purifying selection (conservation), while negative values (red) indicate a significant drive for physicochemical change.

**Figure 1:**
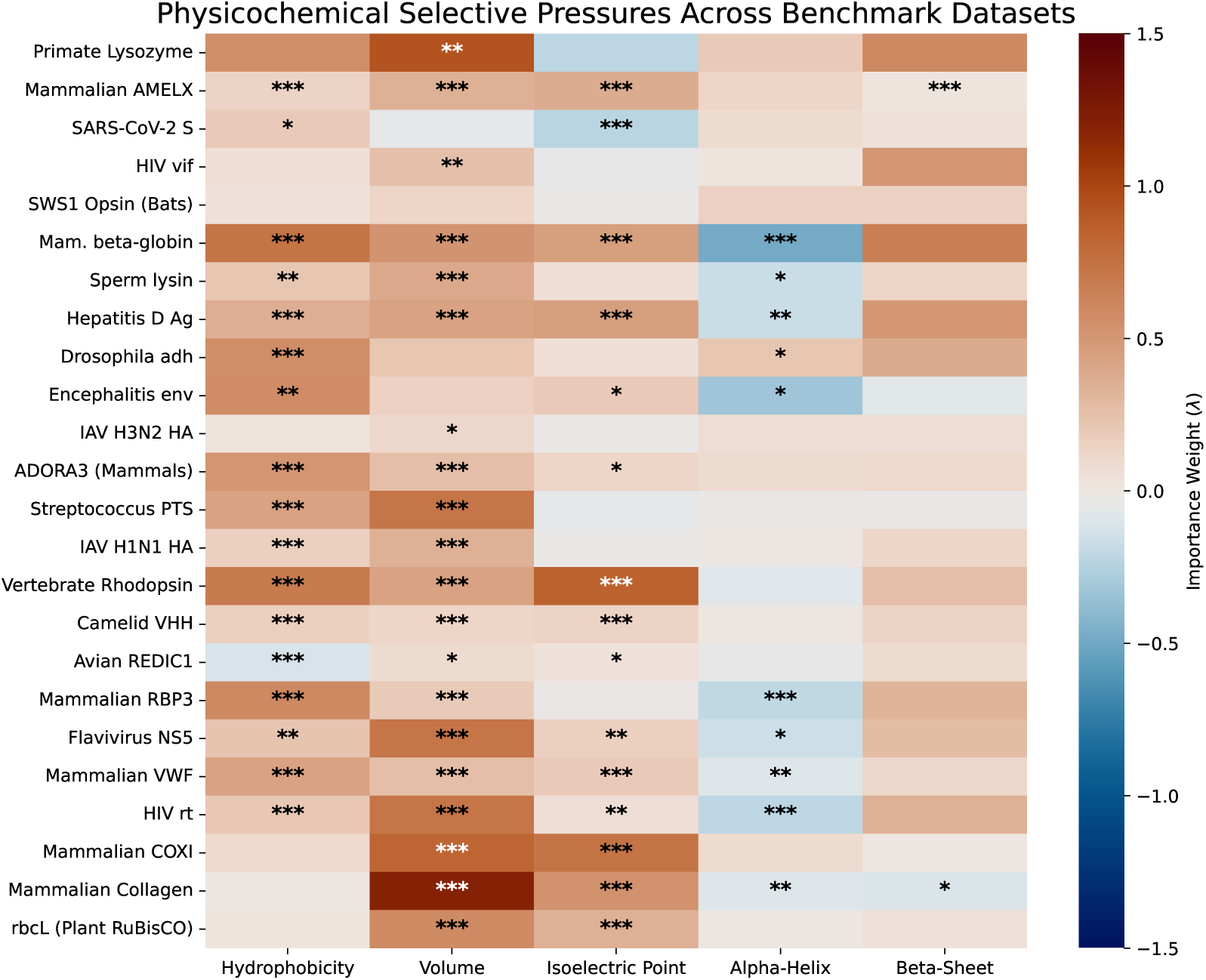
Physicochemical selective fingerprints across 24 benchmark datasets. The heatmap displays estimated property importance weights (*λ*) for the 5-prop PRIME model. Blue cells indicate property conservation (purifying selection), while red cells indicate a drive for property change (diversifying selection). Asterisks denote *λ* values significantly different from zero (Likelihood Ratio Test; * *p <* 0.05, ** *p <* 0.01, *** *p <* 0.001). Datasets are ordered by total evolutionary information (*L* × *T*). Hydrophobicity and volume emerge as nearly universal constraints, while secondary structure propensities and isoelectric point serve as the primary substrates for adaptive tuning.

### 3.4 Capturing Episodic and Physicochemical Heterogeneity

To formally assess whether explicitly modeling variable physicochemical constraints improves our description of protein evolution, we compared E-PRIME against a strict baseline: a 2-component BUSTED model. Our results demonstrate that the standard assumption of uniform amino acid exchangeability across a sequence is fundamentally insufficient. The substantial improvements in model fit observed for E-PRIME over the BUSTED baseline (Table 3)—exceeding 1800 AICc units for rbcL and 1200 AICc units for Mammalian Collagen—underscore that selective pressures fluctuate not only in magnitude (*dN/dS*) but in the specific biophysical rules governing substitutions.

**Table 3:**
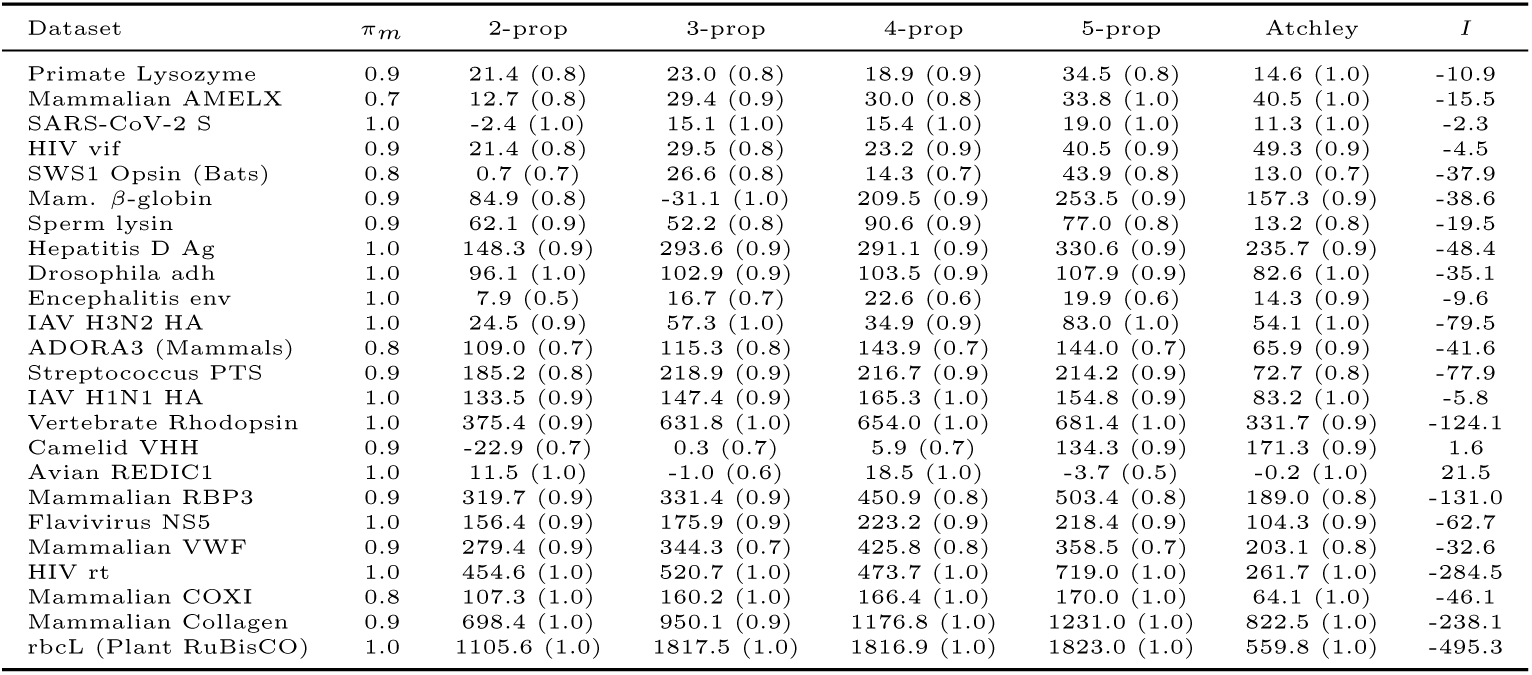
Comparison of E-PRIME models against the BUSTED baseline for 24 benchmark alignments. We report ΔAICc relative to BUSTED, with the weight of the dominant mixture component (*πm*) provided in parentheses for E-PRIME models and in the second column for BUSTED. Datasets are sorted by *L* × *T* . The Interaction Information (*I*) column quantifies synergy (*I <* 0) using the 5-prop model. Best-fitting models (ΔAICc *>* 0) are highlighted in bold.

A compelling example is Vertebrate Rhodopsin, which illustrates the critical importance of modeling specific physicochemical properties. Even the 2-prop E-PRIME model (incorporating only hydrophobicity and volume) yields a large improvement over BUSTED (ΔAICc ≈ 375), confirming that basic structural constraints constitute a primary driver of rate heterogeneity. However, the addition of isoelectric point in the 3-prop model triggers a further substantial jump in fit (ΔAICc ≈ 632). This result reveals that capturing the precise electrochemical constraints required for specialized functions, such as spectral tuning, is essential for a biologically accurate model of evolution. Other datasets reinforce this mechanistically grounded logic. Hepatitis D Antigen exhibits a large gain in fit with the 3-prop model (ΔAICc_3_*_−_*_2_ ≈ 145), consistent with its role as an RNA-binding protein where surface electrostatics mediate interactions with the phosphate backbone. Similarly, Mammalian Collagen shows a marked fit improvement with E-PRIME 5-prop (ΔAICc ≈ 1231), further validating the dominance of volume constraints in its rigid triple helix.

The mixture components recovered by E-PRIME (Supplementary Table S4) reveal a pervasive biophysical dichotomy between structural preservation and functional adaptation. The dominant regime, typically assigned a weight above 70%, is characterized by intense conservation of hydrophobicity and volume, likely representing the requirement to maintain the buried protein core. In sharp contrast, the minor regime captures the distinct biological modes of each protein. In viral surface antigens such as SARS-CoV-2 Spike or IAV H3N2, this component identifies diversifying selection acting on charge or hydrophobicity, potentially reflecting active immune escape. In metabolic enzymes like rbcL or HIV-1 rt, the minor regime represents a more permissive purifying environment where constraints are relaxed but directional change remains rare.

Finally, we interrogated whether the improvements offered by rate heterogeneity (BUSTED) and physicochemical heterogeneity (PRIME) are redundant or synergistic. We quantified this relationship by calculating the Interaction Information (*I*), an information-theoretic metric defined as *I* = ΔAICc_PRIME_ +ΔAICc_BUSTED_ −ΔAICc_E-PRIME_, where ΔAICc represents the improvement over the MG94 baseline. We found that for the vast majority of datasets—including rbcL (*I* ≈ −495 AICc) and Mammalian Collagen (*I* ≈ −238 AICc)—the interaction was strongly synergistic (*I <* 0). While *I* serves as a useful proxy for synergy by comparing independent model fits, we emphasize that it does not replace a formal joint hierarchical analysis. Nevertheless, the pervasive negative values suggest that rate and property constraints capture orthogonal dimensions of the evolutionary process: a site is often conserved in rate precisely *because* its substitutions are filtered through a specific biophysical sieve. Modeling both dimensions simultaneously allows for a more precise resolution of these selective forces.

### 3.5 The Landscape of Physicochemical Constraints

To evaluate the broader applicability of our framework, we deployed PRIME and E-PRIME to characterize the physicochemical landscape of the mammalian proteome. We analyzed a high-quality subset of 18,944 mammalian genes from a larger collection of approximately 19,000 alignments. This subset was selected based on several criteria to ensure robust inference: alignments were required to contain at least 10 sequences, a length of at least 50 codons, and a total tree length between 0.5 and 50 substitutions per site. Furthermore, we only included genes where successful fits were obtained for all five comparison models (MG94, CoRa, PRIME, E-PRIME, and BUSTED).

Given that the 5-property model provided the optimal fit for the majority of benchmark datasets (Tables S2 and 3) and successfully captured critical structural constraints missed by simpler parameterizations, we restricted this large-scale analysis to the G-PRIME and E-PRIME 5-prop models. This approach allows for a computationally efficient yet biologically comprehensive characterization of selective pressures across the proteome. An aggregate, gene-wide model of physicochemical evolution should effectively serve as a low-pass filter, resolving the dominant selective pressures that shape the majority of residues.

We formulated three primary biophysical expectations for this genome-wide model. First, the Baseline Expectation posits the pervasive conservation of hydrophobicity and volume, as maintaining the hydrophobic core and preventing steric clashes are universal requirements for protein folding and solubility (Dill, 1990; Richards, 1974). Second, we anticipated a Structural Signal where beta-sheet propensity is more strictly conserved than alpha-helix propensity, reflecting the rigid, cooperative nature of beta-sheet scaffolds. Third, any Regulatory Mode involving diversifying selection (*λ <* 0) or neutrality should primarily target alpha-helix propensity, consistent with the prevalence of Intrinsically Disordered Regions (IDRs) and flexible linkers where helical stability is tuned to modulate binding affinity (Zarin et al., 2019).

Our analysis of 18,944 mammalian genes provides a compelling validation of these biophysical principles. Consistent with the Baseline Expectation, we observed a pervasive signature of purifying selection acting on core properties. As illustrated in the distributions (Figure 2B), Hydrophobicity (*λ̅* ≈ 0.265) and Volume (*λ̅* ≈ 0.301) predominantly exhibit positive importance weights. This conservation is not merely numerical but statistically robust; Likelihood Ratio Tests (LRTs) confirm that constraints on residue volume and hydrophobicity are significant (FDR ≤ 0.05) in 85.8% and 74.2% of the proteome, respectively. Furthermore, confirming our Structural Signal, we find that Beta-Sheet propensity is subject to widespread conservation (*λ̅* *>* 0; significant in 75.5% of genes). In sharp contrast, and validating the Regulatory Mode expectation, Alpha-Helix propensity exhibits a uniquely broad distribution centered near zero (*λ̅* ≈ −0.092). While helical structure is conserved in many contexts (16.0%), a substantial fraction of the proteome shows either neutrality or active selection for change (44.0%, *λ <* 0), identifying alpha-helix propensity as a primary axis of evolutionary tuning.

**Figure 2:**
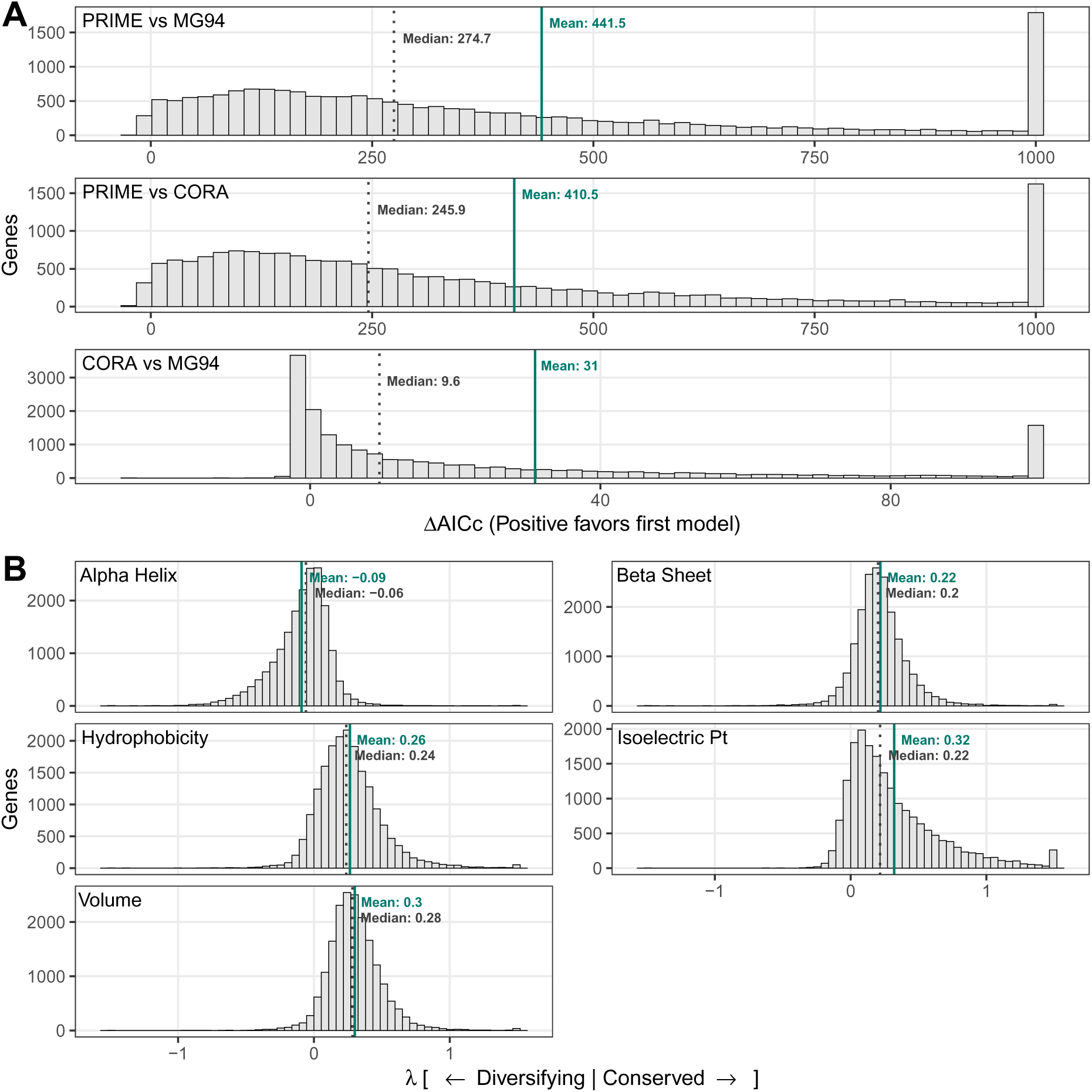
Genome-wide analysis of physicochemical selection in 18,944 mammalian genes. (A) Model selection performance showing distributions of ΔAICc for three comparisons: Global 5-prop PRIME vs. MG94 baseline, PRIME vs. CoRa, and CoRa vs. MG94. Histograms are faceted by comparison with independent scales; values are clamped at -25 and +1000 for visibility. (B) Distributions of global property importance weights (*λ*) across the proteome. Positive *λ* values signify property conservation (purifying selection), while negative values signify a drive for physicochemical change (diversifying selection). Data are clamped at ±1.5 for visualization. Summary statistics (mean, median, and 95% intervals) for each distribution are provided in the inset boxes.

To dissect the combinatorial logic of these constraints, we discretized property weights into conserved, changing, or neutral states (FDR ≤ 0.05). Analysis of the top 20 most frequent patterns (Figure 3) reveals that the predominant mode of mammalian evolution is indeed pervasive conservation across most dimensions. Selection for change is remarkably sparse and targeted; in the rare instances where a property is under significant drive to change, it is almost exclusively restricted to a single property: alpha-helix propensity. For example, the most common regime (23.5% of genes) corresponds to the simultaneous conservation of hydrophobicity, volume, isoelectric point, and beta-sheet propensity, coupled with significant selection for changing alpha-helix propensity. Collectively, these results underscore that while core packing properties are almost universally rigid, the evolutionary tuning of secondary structure—particularly alpha-helices—constitutes a major axis of mammalian adaptation.

**Figure 3:**
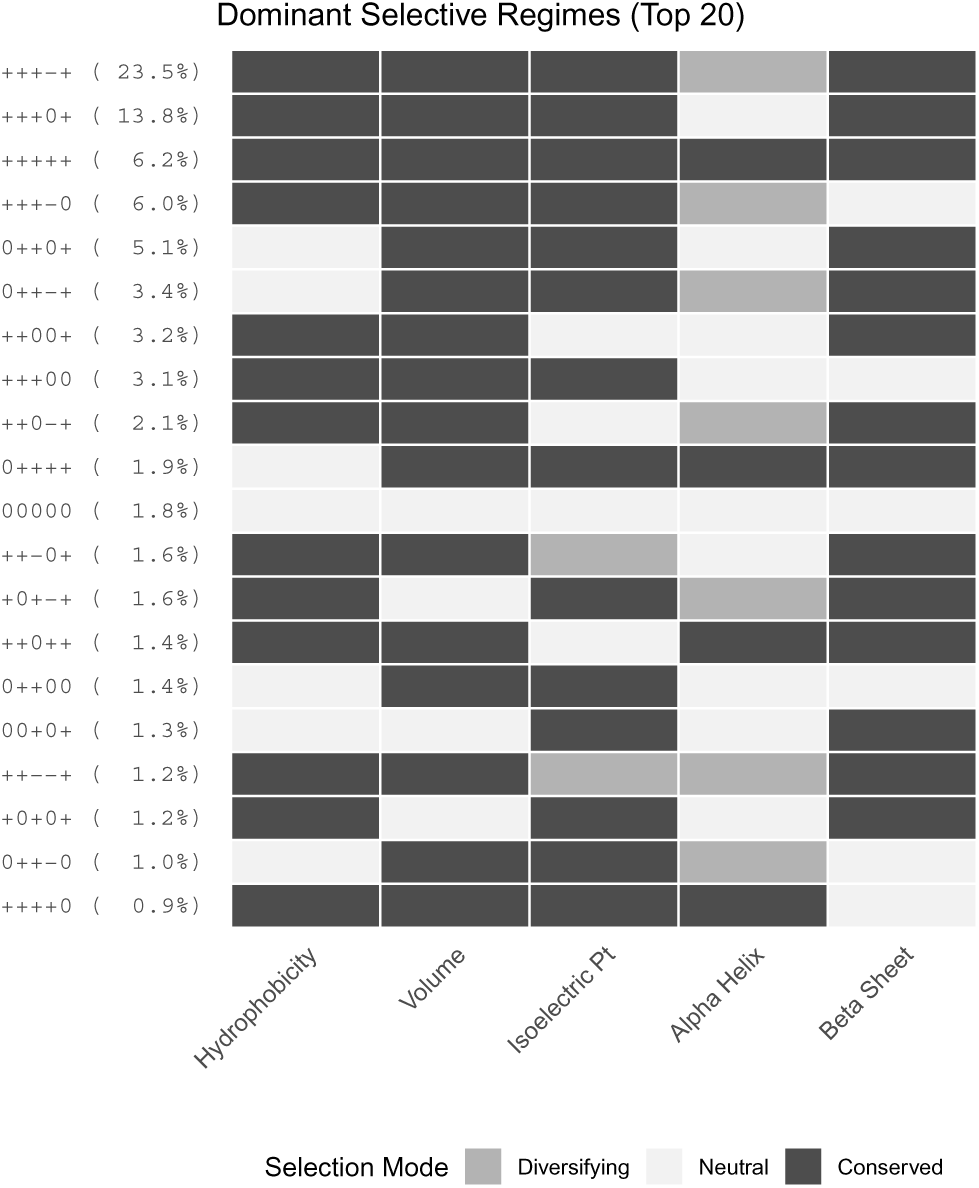
Dominant Physicochemical Selective Regimes. The top 20 most frequent combinations of property constraints across 18,944 mammalian genes. Properties are ordered as: Hydrophobicity, Volume, Isoelectric Point, Alpha-Helix, Beta-Sheet. For each property, genes were classified as Conserved (dark gray), Changing (light gray), or Neutral (white) based on an FDR threshold of 0.05. The patterns are ordered by frequency, with the percentage of the proteome shown in parentheses. The most common regime involves broad conservation coupled with selection for changing alpha-helix propensity.

While the dominant trend is the singular diversification of alpha-helix propensity, multi-property diversification regimes do exist and point to specific evolutionary conflicts. For instance, the + + − − + regime (1.2% of genes), which combines the conservation of core structure (hydrophobicity, volume, beta-sheet) with simultaneous selection for changing isoelectric point and alpha-helix propensity, is strongly enriched for host defense enzymes such as Serine Peptidases (GO:0008236, *p_adj_* = 1.6 × 10*^−^*^11^; e.g., *KLK8*, *ELANE*) and glycosyltransferases (GO:0008417, *p_adj_* = 6.5 × 10*^−^*^6^; e.g., *FUT3*, *FUT6*). This dual diversification likely reflects the need to rapidly adapt surface electrostatics and loop dynamics to engage diverse pathogen substrates while maintaining the catalytic fold.

### 3.6 Case Study: Dissecting Regulatory and Environmental Constraints

To further validate that PRIME captures biologically meaningful selective signals, we performed two case studies contrasting protein families with divergent structural and functional paradigms. First, we compared Cadherins (*N* = 92), which function as rigid extracellular adhesion scaffolds, against Nuclear Receptors (*N* = 46), which act as dynamic intracellular regulatory switches (Figure 4A). While both families exhibit comparable conservation of beta-sheet propensity (*λ̅* ≈ 0.28), they diverge starkly in their evolutionary treatment of alpha-helices. Notably, 76% of Nuclear Receptors show statistically significant diversifying selection on alpha-helix propensity (*λ <* 0), consistent with the fine-tuning of disordered transactivation domains. In contrast, Cadherins show a mixed regime of weak conservation or neutrality, with virtually no systematic drive for diversification (*λ̅* ≈ 0.03).

**Figure 4:**
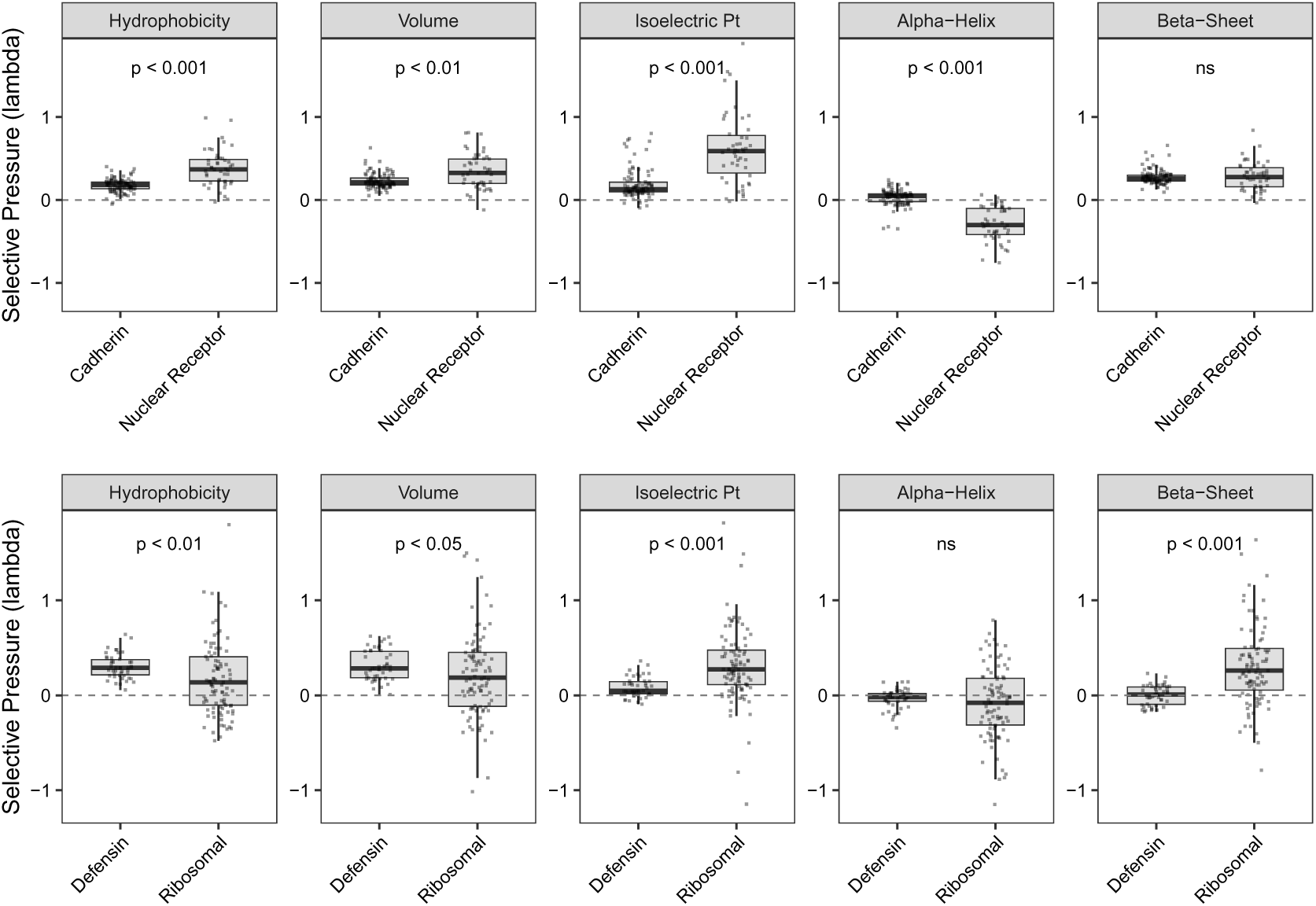
G-PRIME distinguishes structural, regulatory, and interaction-driven constraints. (A) Comparison of global property impor-tance weights (*λ*) between Cadherins and Nuclear Receptors. While both families conserve beta-sheet propensity (structural stability), Nuclear Receptors show strong diversifying selection on alpha-helix propensity (regulatory tuning), contrasting with the neutral evolution in Cadherins. (B) Comparison between Defensins and Ribosomal proteins. Ribosomal proteins exhibit strict conservation of isoelectric point and beta-sheet propensity (RNA-binding and machine assembly), while Defensins show significantly reduced constraint on these properties, consistent with rapid adaptive evolution in immune defense. P-values derived from Mann-Whitney U test are shown above each comparison.

Second, we evaluated interaction-driven constraints by comparing Defensins (*N* = 45) against Ribosomal Proteins (*N* = 95) (Figure 4B). As predicted, Ribosomal proteins exhibit strict conservation of isoelectric point (*λ̅* ≈ 0.41; significant in 54% of genes), reflecting the requirement for positive charge in RNA binding. In sharp contrast, Defensins show significantly lower constraint (*λ̅* ≈ 0.08), with more than half the family (53%) evolving neutrally with respect to charge. This relaxed sieve supports an evolutionary model where surface electrostatics are permitted to drift or adapt, facilitating the recognition of diverse and evolving pathogen surfaces.

### 3.7 Model Selection and Synergy

The 5-property PRIME model consistently outperforms alternative parameterizations across the proteome. While the CoRa model improves upon the MG94 baseline in 76.5% of datasets, G-PRIME (5-prop) offers a far more substantial advance, providing a superior fit for 98.6% of analyzed genes (Figure 2A). Furthermore, PRIME outperformed CoRa itself in 98.5% of cases. This strong support confirms that the site-specific, quantitative modeling of physico-chemical constraints provides a far more accurate description of the evolutionary process than rate-based or binary classification approaches. We further quantified the relationship between rate heterogeneity (captured by BUSTED) and physicochemical heterogeneity (captured by PRIME) by computing the Interaction Information (*I*) across the genome, as defined in Section 3.4. While the addition of rate heterogeneity alone (BUSTED vs. MG94) yielded a mean improvement of 211.1 AICc units, and physicochemical heterogeneity alone (PRIME vs. MG94) yielded 441.5 units, the combined E-PRIME model provided a mean gain of 723.6 units over the MG94 baseline. Consistent with our benchmark findings, we observed pervasive synergy between these dimensions: the mean interaction information was -71.0, with 84.4% of genes exhibiting synergy (*I <* 0). This result demonstrates that for the vast majority of the mammalian proteome, accounting for both temporal rate variation and site-specific physicochemical constraints yields a model fit superior to the sum of their individual contributions, confirming that these frameworks capture orthogonal and synergistic components of the evolutionary process.

### 3.8 The Dynamic Architecture of Physicochemical Constraints

To further explore the nature of this synergy, we analyzed the parameter estimates from the E-PRIME model to map the dynamic architecture of constraints (Figure 5). We found that physicochemical selective pressures are rarely static; the average gene exhibits a secondary evolutionary regime accounting for approximately 18.5% of the phylogenetic history. This analysis reveals a clear hierarchy of constraint. Fundamental packing requirements—hydrophobicity and residue volume—remain comparatively rigid, with fewer than 3% of genes showing significant fluctuations between conserved and diversifying modes. In contrast, surface electrostatics and secondary structure elements serve as the primary substrates for episodic adaptation, exhibiting high plasticity between the majority and minority regimes.

**Figure 5:**
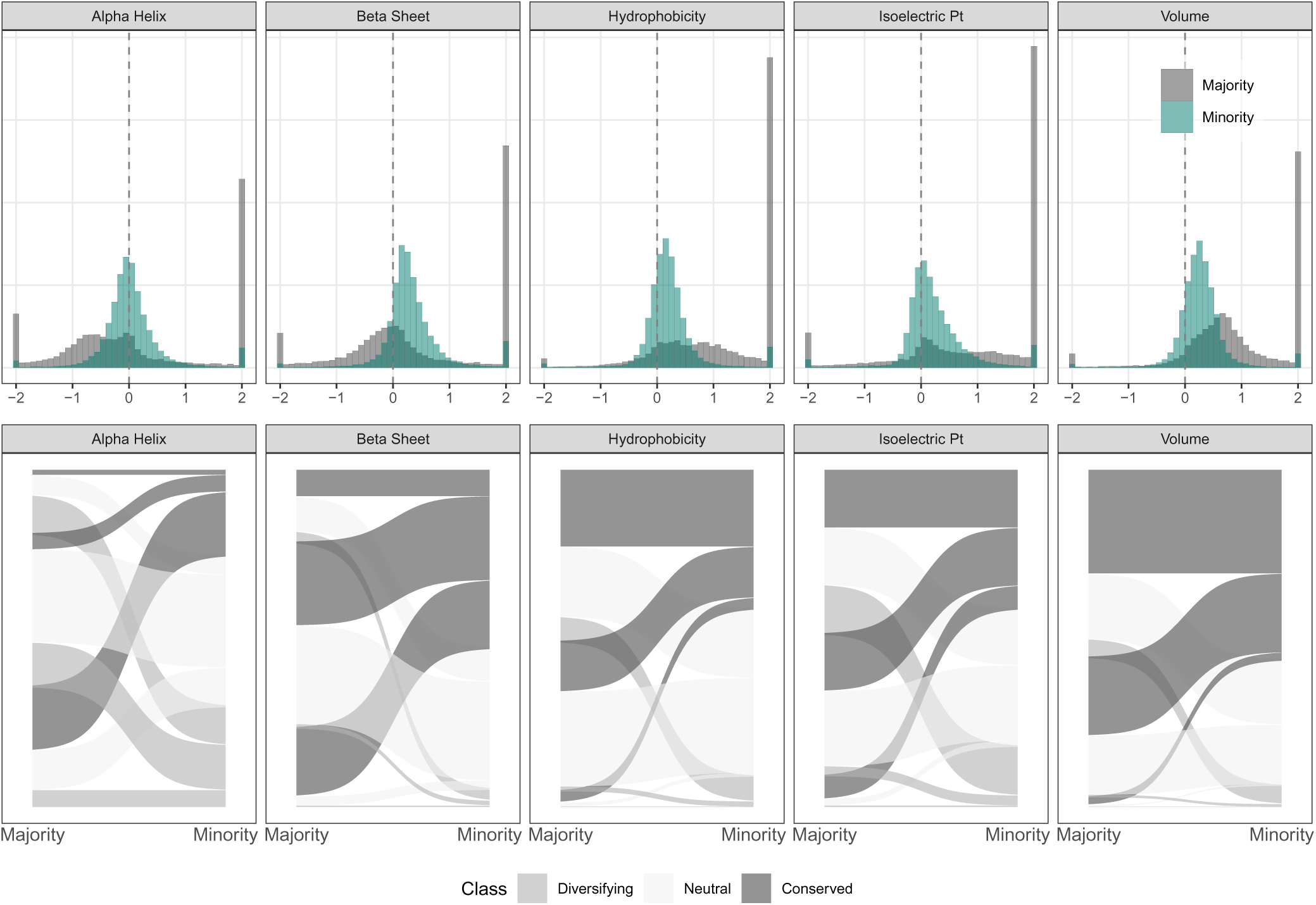
Divergent Selective Regimes in the Mammalian Proteome. (A) Density distributions of property importance weights (*λ*) for the Majority (dark grey) and Minority (teal) mixture components of the E-PRIME model. The Minority component is characterized by intense conservation of Beta-sheet propensity, Volume, and Hydrophobicity, while Isoelectric Point and Alpha-Helix propensity show greater variance. (B) Alluvial plot tracking the flow of evolutionary states (Conserved, Neutral, Diversifying) from the Majority to the Minority component (FDR ≤ 0.05). Note the dramatic shift in Beta-sheet propensity from neutral/diversifying in the Majority to conserved in the Minority (scaffold regime), and the increased diversifying selection on Isoelectric Point in the Minority regime.

A notable feature of this landscape is the scaffold signature, characterized by diversifying selection on Beta-sheet propensity in the majority component but strict conservation in the minority (*λ_maj_ <* 0*, λ_min_ >* 0). This pattern, observed in 20.3% of genes, suggests that for many proteins, maintaining specific beta-strand elements is critical even when the global fold favors alternative structures. The Pleckstrin Homology domain-containing family G (PLEKHG) serves as a paradigm for this regime (86% of members), likely reflecting the strict structural conservation of their beta-rich PH domains amidst diversifying regulatory regions. Conversely, the charge tuning signature (*λ_maj_ >* 0*, λ_min_ <* 0) defines a mode of functional adaptation where global electrostatic stability is maintained while specific interfaces are driven to change. This regime is significantly enriched for Olfactory Receptors (GO:0004984, *p_adj_ <* 10*^−^*^20^), enabling the rapid exploration of chemical space required for chemosensation. Analysis of gene families reinforces this finding: the Fucosyltransferase (FUT) family (64% of members) and NLRP innate immune sensors (57%) both strongly exhibit this signature, consistent with the adaptive retuning of recognition surfaces in host-pathogen co-evolution.

Finally, we identified rare but biologically coherent regimes associated with intense evolutionary conflict. The double diversification signature for isoelectric point (*λ_maj_ <* 0*, λ_min_ <* 0) implies pervasive selection for charge alteration across the entire protein and is enriched in antiviral factors such as *APOBEC3* (30%) and *SLFN* (67%).

Similarly, cryptic diversification (*λ_maj_* ≈ 0*, λ_min_ <* 0) is a hallmark of immune receptor families like *KIR* (67%) and *LILRA* (80%), as well as antimicrobial peptides such as *Defensins* and *Cathelicidins* (GO:0019730, *p_adj_* ≈ 4 × 10*^−^*^5^). This suggests that for recognition factors, the focused adaptation of electrostatic binding interfaces is the primary evolutionary mode, occurring even when the majority of the structure remains evolutionarily neutral.

Finally, we examined genes exhibiting discordant selection on packing constraints (e.g., *λ_Hydro_*conserved in one component but diversifying in the other). This regime is significantly enriched for transcription factors (GO:0006357, *p_adj_* ≈ 0.01), particularly Zinc Finger proteins. This likely reflects the modular architecture of these proteins, which alternate between rigidly packed DNA-binding domains (conserved core) and flexible, disordered transactivation regions (relaxed or diversifying packing). These results suggest a hierarchical architecture of constraint: while the packing of the hydrophobic core remains non-negotiable, surface electrostatics and secondary structure elements serve as the primary substrates for episodic adaptation.

### 3.9 Resolving the Biophysical Mechanism of Selection

Aggregate gene-wide models, while useful, average over the heterogeneous biophysical requirements of individual residues. Here, we deploy the S-PRIME inference framework to resolve evolutionary forces at single-residue resolution. By estimating a vector of property weights (*λ*) for each site, we move beyond the phenomenological detection of selection to explicitly identify *which* physicochemical properties are constrained or diversified. We apply this high-resolution lens to (1) uncover cryptic constraints in sites that appear neutral by standard rate-based metrics, (2) map the structural anatomy of physicochemical pressure, and (3) dissect the biophysical logic of key adaptations in viral and mammalian proteins.

### 3.10 Motivating Example: Selection in Influenza Hemagglutinin

To illustrate how PRIME partitions evolutionary signals into specific biophysical mechanisms, we analyzed the evolution of Influenza A H3N2 Hemagglutinin (HA). As the primary surface antigen, HA is subject to intense selective pressure to evade host immunity while maintaining receptor binding and membrane fusion. Standard methods (*dN/dS*) effectively identify sites under positive selection (Yang et al., 2000a; Murrell et al., 2012b), such as those within the canonical antigenic epitopes. However, these metrics remain agnostic to the biophysical constraints that channel these adaptive shifts. Consider Site 226, located within the receptor binding pocket of the HA head domain. In our H3N2 benchmark dataset, this site exhibits a clear signature of diversifying selection (*dN/dS* ≈ 22, FEL *p <* 10*^−^*^10^). This rate elevation is driven by Glutamine-to-Leucine (Q226L) substitutions, which shift viral specificity from avian-type to human-type sialic acids (Rogers and Paulson, 1983). While *dN/dS* correctly flags this position as rapidly evolving, it treats the Q : L substitution as an abstract token exchange.

In contrast, S-PRIME analysis explicitly resolves the biophysical program of this functional switch, classifying it as an instance of constrained diversification. The model identifies significant diversifying selection on Hydrophobicity (*λ* = −2.49*, p_adj_ <* 10*^−^*^11^) and Volume (*λ* = −1.81*, p_adj_ <* 10*^−^*^5^), capturing the chemical transition from polar Glutamine to a larger, hydrophobic Leucine. This adaptive shift occurs against a backdrop of strict conservation for Isoelectric Point (*λ* = 15.0*, p_adj_* = 0.0004) and Alpha-Helix Propensity (*λ* = 11.8*, p_adj_* = 0.001). Structural mapping (PDB: 4FNK) reveals the biophysical basis for this complexity: Site 226 is largely buried within the receptor binding pocket (RSA = 0.06) and forms part of a beta-bridge. This environment necessitates strict preservation of the local electrostatic and packing architecture, channeling adaptation into a specific chemical trajectory rather than permitting random variation.

The utility of this framework is further underscored by Site 145 (Epitope A). Located in an intermediate solvent-accessible environment (RSA = 0.14, Beta-strand), the PRIME omnibus test identifies pronounced property-dependence (*Q* = 3.6 × 10*^−^*^11^), yet under strict Holm-Bonferroni correction, no single dimension emerges as the sole driver. This identifies Site 145 as a complex driver, where selection likely acts on a distributed combination of surface properties rather than a single axis. This distinguishes it from sites like Site 186 (Epitope B), which appears neutral by rate metrics (*p*_MEME_ = 0.06) but which PRIME resolves as a cryptic constraint driven by the conservation of Isoelectric Point (*λ* = 13.3*, p_adj_ <* 0.001). By partitioning selection into these interpretable channels, PRIME achieves a level of mechanistic resolution that is fundamentally unattainable with standard codon models.

Across the entire H3N2 hemagglutinin gene (*N* = 329), PRIME identified 12 sites with significant physicochemical constraints (FDR ≤ 0.10). These results provide a complementary view of adaptation: only 5 sites were flagged by both PRIME and standard rate-based methods (MEME, *p <* 0.05). PRIME exclusively identified 7 sites missed by MEME—including Site 186 and Site 310 (Stem)—revealing cryptic biophysical signals that rate metrics overlook. Conversely, MEME identified 12 sites not detected by PRIME, which often represent cases of physicochemically unconstrained diversification. Of these 12 sites, 42% are fully solvent-exposed, including Antigenic Site A positions 133 and 135. Structural analysis confirms that these unconstrained sites lack secondary structure (Coil), allowing immune evasion to proceed through a permissive range of substitutions without disrupting the protein fold. This classification framework is formalized in Figure 6.

**Figure 6:**
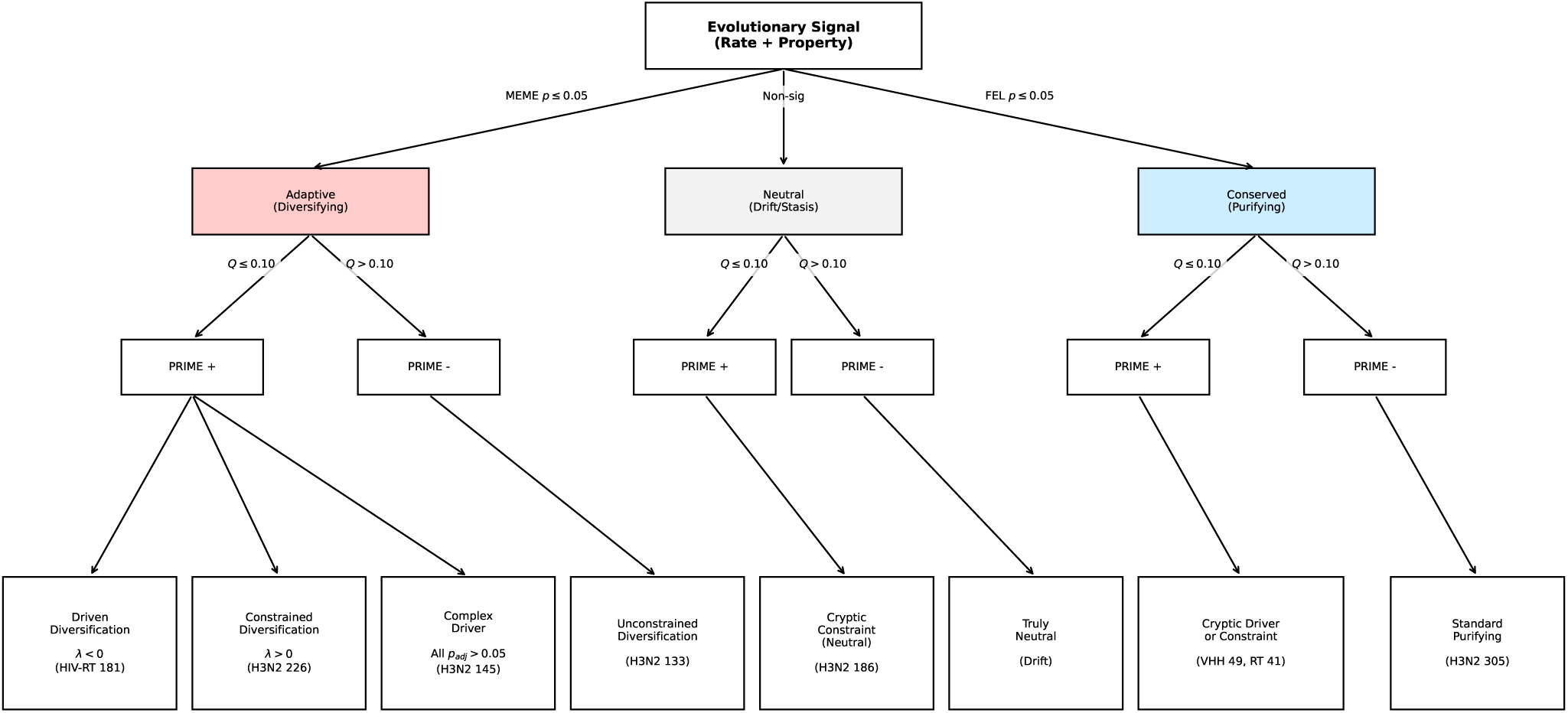
Site Classification Decision Tree. A framework for interpreting site-specific evolutionary signals by integrating substitution rate ratios (*dN/dS*) with physicochemical property weights (*λ*). This approach distinguishes between unconstrained diversifying selection and targeted biophysical adaptation, while also identifying cryptic constraints in neutral (*dN/dS* ≈ 1) or purifying sites. Note: A site may be identified as significant by the PRIME omnibus test (indicating property dependence) even if no single property exhibits a strong individual weight, typically reflecting complex or weak constraints.

### 3.11 The Spectrum of Physicochemical Constraints

By applying our classification framework (Figure 6) across the complete benchmark (*N* = 9, 998 sites), we resolved 576 sites (5.8%) with significant biophysical structure (PRIME FDR ≤ 0.10). Among residues identified as being under diversifying selection (MEME *p* ≤ 0.05), nearly half (45.9%; 180/392) exhibited detectable physicochemical structure. Of these structured adaptive sites, 58.9% resolved as constrained diversification, where adaptation occurs against a backdrop of significant conservation for at least one property. The remaining adaptive sites were classified as Complex Drivers (28.3%) or driven diversification (12.8%), where selection favors property change without detectable structural rigidities. Within the neutral class, 4.3% of sites (226/5,300) exhibited significant structure, identifying them as Cryptic Constraints. Finally, of the 4,306 sites identified as evolving under purifying selection, significant physicochemical constraints were resolved for 170 (3.9%). While the majority of these represent Cryptic Constraints (strict property conservation), we also identify instances of Cryptic Drivers (0.7% of all sites), where specific properties are driven to change despite global purifying selection. Supplementary Figure S1 visualizes these conditional probabilities, revealing the distinct biophysical fingerprints found across the selection spectrum. Detailed counts for each classification category across all benchmark datasets are provided in Supplementary Table S8.

### 3.12 Biological Case Studies in HIV-1 Reverse Transcriptase

To illustrate the biophysical interpretation of S-PRIME parameters, we examined significant sites in HIV-1 Reverse Transcriptase (RT), a protein subject to intense selective pressure from antiretroviral therapy.

Hydrophobicity Conservation (Site 184): Residue 184, located in the conserved YMDD motif of the catalytic site, is a primary determinant of resistance to lamivudine (3TC) and emtricitabine (FTC). S-PRIME identifies this site as undergoing intense purifying selection on hydrophobicity (*λ*_Hydro_ = 15.0, *q <* 10*^−^*^6^) despite a high substitution rate (46 events). The observed mutations (M→V, M→I) involve minimal change in hydrophobicity, preserving the hydrophobic core of the active site while conferring drug resistance. The model correctly infers that while the residue identity can change, the biophysical property—hydrophobic packing—must be strictly maintained.

Volume Conservation (Site 100): Residue 100 is a key site for non-nucleoside reverse transcriptase inhibitor (NNRTI) resistance. Our analysis recovers a strong signature of volume conservation (*λ*_Vol_ = 15.0, *q <* 0.001). Steric constraints in the NNRTI binding pocket limit the set of viable amino acids to those that maintain specific spatial occupancy, a feature captured by the high *λ*_Vol_ estimate.

Charge Diversification (Site 214): In contrast to the conserved active site, residue 214 exhibits a signature of diversifying selection on isoelectric point (*λ*_pI_ ≈ −10.9, *q* = 0.02). This site tolerates and potentially favors radical changes in charge, consistent with surface-exposed residues where electrostatic remodeling may facilitate immune evasion or solubility adaptation without disrupting the protein fold.

### 3.13 Determinants of PRIME Significance

To explicitly resolve the statistical determinants of physicochemical constraint, we performed a logistic regression analysis on 4,558 variable sites (substitutions *>* 0) to predict the probability of identifying significant structure (Omnibus *q* ≤ 0.10). We modeled this probability as a function of the number of inferred substitutions, the amino acid diversity, and their interaction, while controlling for synonymous (*α*) and non-synonymous (*β*) rates. Our analysis reveals that PRIME significance is not merely a proxy for rapid evolution. While the raw number of substitution events is the strongest positive predictor (Odds Ratio ≈ 49 per SD increase, *p <* 10*^−^*^100^), this effect is modulated by a critical negative interaction with amino acid diversity (Interaction Coef = -2.13, *p <* 10*^−^*^13^). This result underscores a fundamental property of the framework: for a fixed number of evolutionary events, the probability of detecting significant structure *decreases* as the site explores a broader range of the amino acid state space. Furthermore, sites with higher non-synonymous rates (*β*) are significantly *less* likely to be identified as structured (Coef = -0.85, *p* = 0.01) once event counts are controlled for, mathematically distinguishing property-driven selection from generic rate acceleration.

These statistical fingerprints demonstrate that PRIME specifically targets toggling regimes—sites where frequent evolutionary turnover is strictly confined to a small physicochemically constrained subset of amino acid substitutions. For instance, Site 184 in HIV-RT exhibits 46 inferred substitutions yet remains restricted to just 3 amino acids (M, V, I). PRIME correctly identifies this as a highly structured regime (*Q <* 10*^−^*^6^), driven by the strict conservation of hydrophobicity (*λ* = 15.0). In stark contrast, Site 75 in Camelid VHH undergoes 21 substitutions and explores 9 different amino acids. Although standard methods identify this site as undergoing significant diversifying selection (MEME *p* = 0.037), PRIME assigns it a non-significant weight (*Q* = 0.53). Structural analysis confirms this residue is highly solvent-exposed (RSA = 0.58), characterizing it as unconstrained diversification rather than the specific structural toggling observed at Site 184. However, with sufficient evolutionary information (e.g., *>* 50 substitutions), PRIME can recover subtle physicochemical biases even in highly variable sites (e.g., Camelid VHH Site 53, 18 amino acids, *Q* = 0.006), suggesting that weak but persistent structural preferences shape even the most hypervariable regions. Even in regimes of low turnover, PRIME can detect significant constraints if the substitutions are highly specific. For example, HIV-RT Site 151 (a known drug resistance position) undergoes only 4 inferred substitutions but is identified as highly significant (*Q <* 10*^−^*^6^), driven by a specific toggle between two amino acid states that modulates hydrophobicity (*λ* = −4.6) while strictly conserving charge.

Empirically, our benchmark suggests that PRIME effectively resolves constraints when sites exhibit this toggling behavior: for positions restricted to 2 or 3 amino acid states, detection power exceeds 65% with as few as 6–10 substitutions. Conversely, highly variable sites exploring broad chemical space (*>* 8 amino acid states) generally require substantial evolutionary turnover (*>* 20 substitutions) to distinguish specific structural preferences from neutral drift.

### 3.14 Decoding Latent Structural Semantics in protein language models

To investigate whether the explicit physicochemical constraints modeled by S-PRIME capture the same fundamental biological structure as implicit deep learning representations, we compared site-specific property weights (*λ*) against residue embeddings from the ESM-2 protein language model (Lin et al., 2023). For each of the benchmark datasets, we computed per-residue embeddings for the primary reference sequence and performed Principal Component Analysis (PCA) to distill the high-dimensional PLM representation into its dominant axes of variation. We then quantified the alignment between the two frameworks by calculating the Spearman rank correlation (*ρ*) between the S-PRIME importance weights and the top principal components of the aligned ESM-2 latent space.

Confirming this hypothesis, we observed statistically significant, albeit modest, correlations across the benchmark datasets (mean max |*ρ*| ≈ 0.19, max ≈ 0.34). This correspondence is particularly notable given the fundamental differences between the two frameworks: S-PRIME infers constraints from a single alignment using an explicit biophysical model, whereas ESM-2 learns statistical patterns from an extensive universe of unrelated sequences. While the relatively low magnitude of these correlations suggests that ESM-2 captures higher-order contextual features that a marginal site-specific model misses, the fact that PRIME’s properties—specifically hydrophobicity and vol-ume—consistently track the principal axes of the PLM latent space suggests that S-PRIME effectively recovers the core biophysical rules that partially underlie the opaque logic of deep learning. This alignment provides independent, data-driven validation that the physicochemical properties modeled here are indeed primary semantic features governing protein evolution.

### 3.15 The Structural Anatomy of Physicochemical Pressure

Mapping site-specific *λ* values to experimental structures for 21 datasets (*N* = 5, 938 sites) reveals a clear and statistically significant structural anatomy of selective pressure (Figure 7). We utilized Relative Solvent Accessibility (RSA) as a proxy for structural depth, classifying residues into Buried (RSA *<* 0.05), Intermediate (0.05 ≤ RSA *<* 0.25), and Exposed (RSA ≥ 0.25) layers. As illustrated in Figure 7, conservation of Hydrophobicity and Volume is strongly depth-dependent. Kruskal-Wallis tests confirm significant differences across RSA classes for all five properties (Volume: *p* = 5.8 × 10*^−^*^23^, Isoelectric Point: *p* = 8.0 × 10*^−^*^31^, Hydrophobicity: *p* = 1.4 × 10*^−^*^8^). Pairwise comparisons using Mann-Whitney U tests with Bonferroni correction reveal that the buried core exhibits significantly higher constraint than the solvent-exposed surface for Volume (*p_adj_* = 2.8 × 10*^−^*^8^ for Intermediate vs Exposed) and Isoelectric Point (*p_adj_* = 2.1 × 10*^−^*^6^ for Intermediate vs Exposed). This gradient is most pronounced when comparing Buried vs Intermediate residues for charge (*p_adj_* = 1.2 × 10*^−^*^9^), reflecting the destabilizing effect of burying unpaired charges.

**Figure 7:**
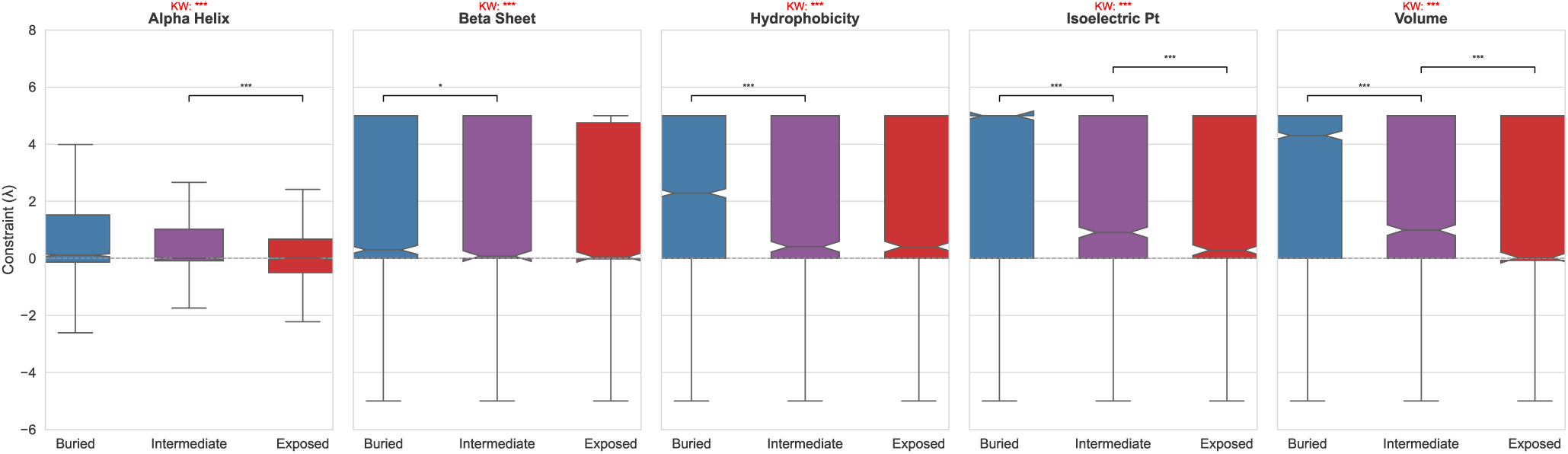
The Structural Anatomy of Selective Pressure. Distributions of site-specific property importance weights (*λ*) stratified by Relative Solvent Accessibility (RSA) classes across 21 benchmark datasets. Properties include (left to right): Alpha-helix propensity, Beta-sheet propensity, Hydrophobicity, Isoelectric Point, and Volume. Brackets indicate significant pairwise differences (Mann-Whitney U test, Bonferroni corrected); Kruskal-Wallis (KW) significance is shown above each panel. Results demonstrate a pervasive gradient where core residues are subject to significantly more stringent physicochemical constraints than surface residues.

While secondary structure propensities also show significant variation (*p <* 10*^−^*^5^), the magnitude of the depth-dependent effect is less severe than for core packing and electrostatics. These results demonstrate that while structural grammar is maintained across the entire fold, the biophysical sieve is most stringent in the protein interior, where packing density and electrostatic neutrality are non-negotiable.

### 3.16 Comparison with Experimental Fitness Landscapes

We compared PRIME inference against Deep Mutational Scanning (DMS) data for the Influenza A H3N2 Hemagglutinin protein. We analyzed 280 variable sites in the A/Perth/16/2009 (H3N2) background, comparing the predicted amino acid preferences from S-PRIME imputation with the experimental fitness measurements. For comparison, we also evaluated zero-shot predictions from the ESM-2 protein language model. S-PRIME shows a moderate correspondence with experimental results, yielding a mean Pearson correlation of *r* = 0.605 and a mean Spearman rank correlation of *ρ* = 0.157 across all variable sites. In comparison, ESM-2 achieved a lower Pearson correlation (*r* = 0.533) but a higher Spearman correlation (*ρ* = 0.305). The mean Jensen-Shannon Divergence (JSD) for S-PRIME was 0.382, while ESM-2 exhibited a lower divergence (*JSD* = 0.327). These results suggest that while global representations from large-scale protein language models better capture the ranking and information-theoretic structure of the fitness landscape, PRIME’s explicit biophysical model provides a competitive linear correspondence and superior mechanistic interpretability.

To further assess the model’s ability to identify the most viable residues, we defined a strongly preferred set for each site (amino acids with predicted or experimental probability *>* 0.10). S-PRIME achieved a mean Jaccard index of 0.501, comparable to ESM-2 (0.460). Furthermore, the residues in the PRIME preferred set capture, on average, 21.2% of the experimental preference mass, a value slightly lower than ESM-2 (21.9%). While this coverage is limited, it demonstrates that the biophysical constraints inferred from evolutionary history align with a meaningful subset of the experimentally determined viable state space, performing comparably to state-of-the-art predictive frameworks. A site-specific dissection of several significant biophysical drivers (Table S9) reveals that PRIME identifies residues where specific properties appear to influence viability. For example, Site 186 (Epitope B), which appears neutral by traditional rate-based metrics, is resolved as a cryptic constraint driven by the conservation of Isoelectric Point (*λ* = 13.3). DMS results show a somewhat restricted state space at this position, with preferences broadly aligning with chemically similar residues (e.g., ASN, GLY, CYS). Similarly, Site 226 exhibits diversifying selection on Hydrophobicity and Volume, reflecting the biophysical transition required for host-receptor adaptation. These patterns are visually corroborated by the side-by-side comparison of S-PRIME, DMS, and ESM-2 logos (Figure 8). Collectively, these results suggest that PRIME recovers aspects of the biophysical rules governing protein fitness, providing a useful surrogate for interpreting experimental landscapes from phylogenetic history.

**Figure 8:**
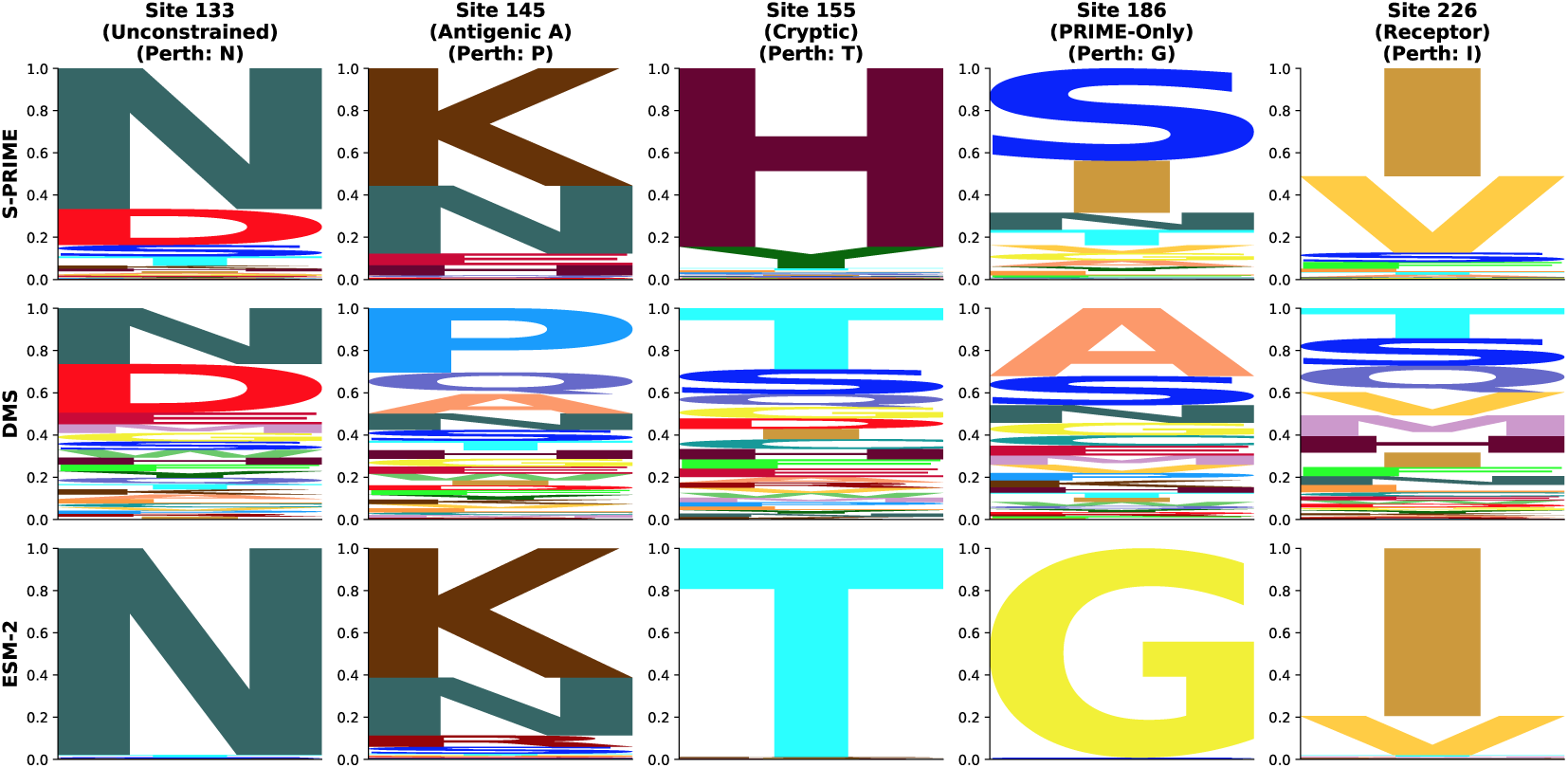
Validation against Experimental Fitness Landscapes. Side-by-side comparison of amino acid preferences for five key sites in H3N2 Hemagglutinin, as determined by S-PRIME phylogenetic imputation (top row), experimental Deep Mutational Scanning (DMS, middle row), and zero-shot predictions from the ESM-2 protein language model (bottom row). Letter height represents relative probability or preference. S-PRIME effectively identifies the subset of chemically viable residues, capturing both strict conservation (e.g., Site 186) and targeted biophysical diversification (e.g., Site 226), performing comparably to the large-scale ESM-2 framework.

### 3.17 S-PRIME Performance Assessment

We assessed the statistical performance of S-PRIME through a comprehensive simulation study encompassing approximately 5,000 replicate alignments across 21 selective scenarios. Analysis of the neutral baseline (scenario null mg94) demonstrates strong calibration of the omnibus test (Supplementary Figure S3A). We find that the test is generally conservative: across 69,774 variable neutral sites, the raw False Positive Rate (FPR) at *α* = 0.05 is only 1.9%, significantly below the nominal threshold. This conservative behavior arises because the 5-property model is over-parameterized for typical sites with low informational depth, leading to a p-value distribution that is stochastically larger than uniform. Following Benjamini-Hochberg correction (*q* ≤ 0.10), the mean FPR across all datasets is effectively 0.00%, confirming that the framework is highly specific and robust to stochastic noise under neutral evolution. This robust error control is further bolstered by the conservative bulk null distribution, which ensures that only the most informative biophysical signals survive multiple testing correction. Detection power is strongly stratified by informational content, with data-rich alignments achieving high sensitivity (Supplementary Figure S3B).

Detection power is primarily governed by the informational redundancy of substitutions at a site rather than global tree length alone (Figure 9C-F). We identify the redundancy ratio (*R* = *N_subs_/N_aa_*) as the fundamental engine of detection, defining the practical boundaries of biophysical discovery. Formal logistic regression across ∼ 200, 000 simulation sites (alternative model) confirms that *R* is a highly effective predictor of detection sensitivity (AUC = 0.91). Power scales sharply and non-linearly with *R*: predicted probability of detection is low (7.9%) at sites with *R* = 1.0, but increases to 28.5% at *R* = 2.0, reaching 49.4% at *R* = 3.0 and exceeding 93.4% when *R* ≥ 10.0 (Figure 9F). Notably, even small datasets can achieve near-perfect sensitivity provided they reach sufficient mutational depth (*R >* 5.0, Figure S2). Our visualization of the informational landscape (Figure 9) identifies distinct regimes of performance. We identify a robust sweet spot for discovery at sites with moderate to high substitution counts (*N_subs_ >* 10) and low to moderate amino acid diversity (*N_aa_*∈ [2, 5]). In this regime, the redundancy ratio exceeds 2.0, providing the mutational depth required to resolve specific property importance weights while minimizing the property-independent noise. Sensitivity in this region consistently exceeds 60%, even under strict FDR control (Panel D).

**Figure 9:**
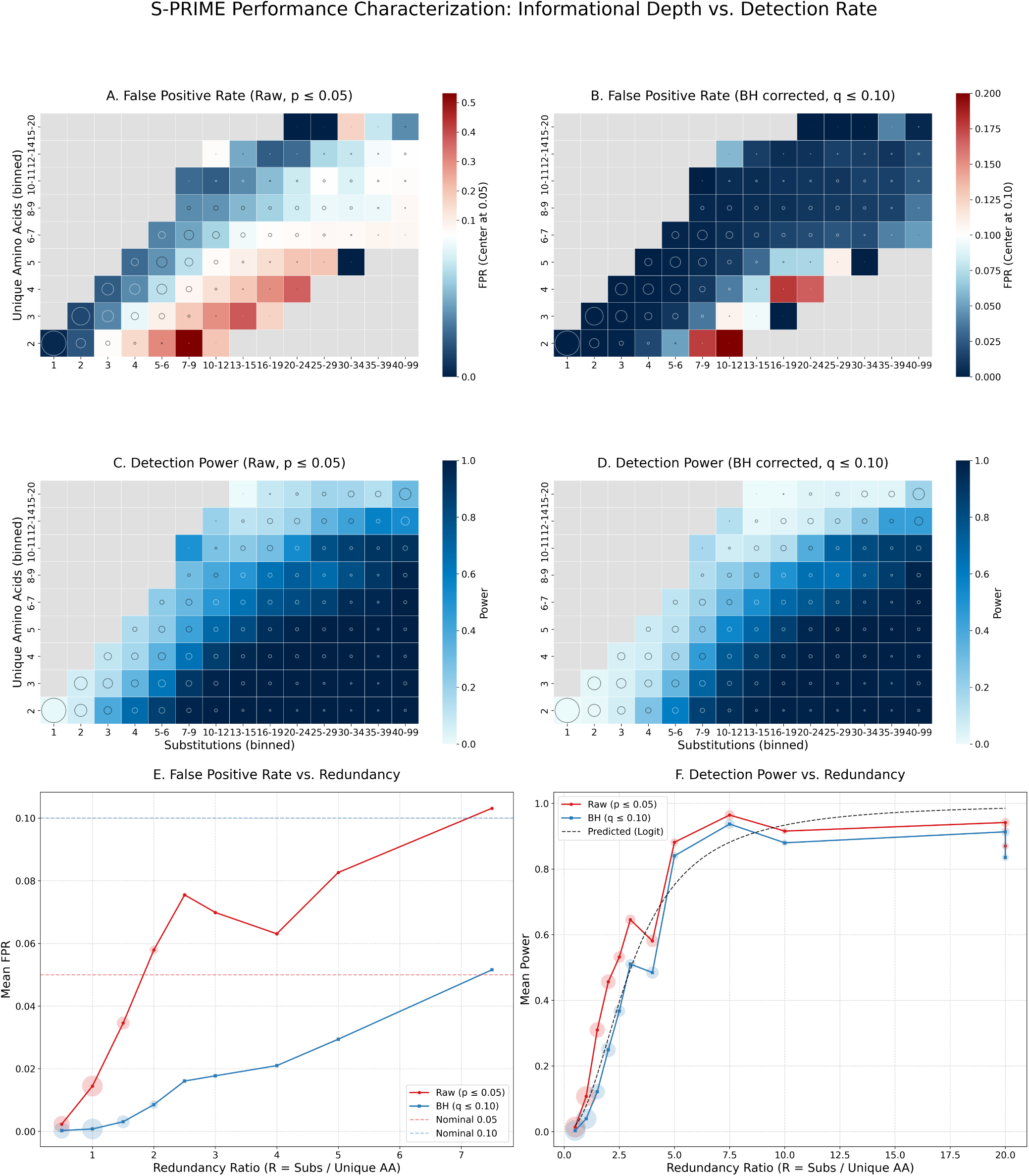
Informational determinants of S-PRIME performance. (A-D) Binned heatmaps illustrating the relationship between the number of non-synonymous substitutions, amino acid diversity, and detection rates. FPR (Panels A, B) is evaluated on the Null partition (*λ* = 0), while Power (Panels C, D) is evaluated on the Alternative partition. Panels A and C show results for the raw omnibus test (*p* ≤ 0.05), while B and D show results after Benjamini-Hochberg correction (*q* ≤ 0.10). Heatmap cells represent mean rates across all replicates; cell area (adaptive bubble strokes) denotes the relative number of simulation sites in that bin. Cells with no simulation data are colored gray. (E, F) Pooled performance as a function of the redundancy ratio (*R*). Bubble area in E and F is proportional to the total number of sites in each *R* bin.

Conversely, we detect three primary danger areas where biophysical signal is either extinguished or misidentified. The first is the invariance trap, located at the informational origin (Figure 9C,D, bottom-left). Sites with few substitutions or extremely rigid constraints (*λ >* 10) lack the requisite variation to characterize the biophysical filter, resulting in 0% power. This simply confirms that S-PRIME is fundamentally a comparative method that requires observed substitutions to quantify the relative costs of residue exchange. The second danger area is the toggling artifact observed at sites with high substitution counts but minimal amino acid diversity (*N_aa_* = 2; Figure 9A). Biologically, repeated toggling between only two residues across deep phylogenies is highly informative of a strict evolutionary constraint. Statistically, however, attempting to map this restricted substitution pathway onto five independent physicochemical parameters creates an under-determined system. Because multiple combinations of the five *λ* weights can perfectly isolate the two observed states from the unobserved chemical space (complete separation), the likelihood optimizer frequently diverges toward parameter boundaries. When maximum likelihood estimates hit these boundaries, the asymptotic assumptions of Wilks’ Theorem break down; the Likelihood Ratio Test statistic no longer follows a standard *χ*^2^ distribution, leading to significantly inflated raw False Positive Rates (Figure 9A). Formal logistic regression on 1,111,446 neutral simulation sites confirms that *R* is a strong predictor of spurious detection (AUC = 0.76), with the predicted raw FPR rising from 1.8% at *R* = 1.0 to 18.3% at *R* = 5.0 and exceeding 65.0% when *R* ≥ 20.0. Fortunately, we observed empirically that the Benjamini-Hochberg correction effectively filters out these mathematically unidentifiable artifacts (Figure 9B). This correction is highly robust because high-redundancy toggling sites are extremely rare under neutral evolution; our analysis of 69,774 simulated neutral sites found that while biallelic sites (*N_aa_*= 2) are common, fewer than 0.1% exhibit the mutational depth (*R >* 2.0) required to trigger severe LRT inflation. Nevertheless, we caution that site-level attribution of specific properties is statistically under-determined for highly restricted sites, and S-PRIME achieves peak identifiability at sites exploring at least 3 distinct amino acid states. The third danger area is the saturation zone (Figure 9C,D, top-right), where sites undergo high levels of turnover across a broad range of amino acids (*N_aa_ >* 10). In these hypervariable regimes, the biophysical signal is so diluted by high property-independent noise, that it requires a very high number of susbtitutions to resolve (Figure 9F).

Stratifying performance by individual datasets and selective regimes (Supplementary Figure S3) reveals a strong dependence of average detection power on informational content. We find that sensitivity is primarily a function of the total mutational depth (*L* × *T*); large, divergent alignments such as *rbcL* (RBCL), Mammalian Collagen (COLL), and HIV-1 RT (HVRT) achieve near-universal power across nearly all selective scenarios (Panel B). In contrast, sparsely sampled or low-divergence genes like Primate Lysozyme (LYSZ) and Mammalian AMELX (AMEL) remain largely underpowered even under intense selective regimes. This result confirms that the practical utility of S-PRIME is fundamentally constrained by the evolutionary information present in the multiple sequence alignment.

Crucially, the framework exhibits robust error control across this entire diversity of data regimes. As illustrated in the actual FDR heatmap (Figure 9 A), the empirical False Discovery Rate remains tightly calibrated to or below the nominal 0.10 threshold for virtually all benchmark genes. The few exceptions are for datasets that are short and diverse (e.g, Camelid diversifying 5) or large and saturated (e.g, HIV-RT diversifying 8). The scenario-wide averages (final row) confirm that the Benjamini-Hochberg correction effectively maintains a stable and conservative error profile regardless of the underlying biophysical signal, even as the underlying relevance ratio is increased (Figure 9E). Collectively, these results establish S-PRIME as a reliable and statistically sound tool for biophysical characterization, provided that the input datasets meet the informational requirements identified in our informational landscape analysis.

## 4 Discussion

In this study, we introduced the PRIME framework, a suite of codon-based evolutionary models that incorporate biophysical properties to characterize the mechanistic basis of natural selection. By applying these models across a taxonomically diverse benchmark and a genome-wide screen of nearly 19,000 mammalian genes, we identified the expected fundamental hierarchy of selective constraint: while core packing properties and beta-sheet scaffolds are rigidly conserved to maintain structural integrity, alpha-helix propensity and surface electrostatics serve as the primary substrates for evolutionary adaptation and episodic tuning. At the site-specific level, S-PRIME resolved the biophysical basis of key functional switches and identified constraints in residues that appear neutral by traditional *dN/dS*-based metrics. Furthermore, our validation against both experimental fitness landscapes and protein language models demonstrates that explicit physicochemical features effectively capture the biophysical principles of selection, providing an interpretable alternative to predictive frameworks that lack explicit mechanistic parameters.

The fundamental contribution of PRIME is its ability to resolve the specific biophysical constraints of natural selection, a goal that traces back to the 1970s (Grantham, 1974) but has historically been overshadowed by the intuitive simplicity of biochemically agnostic *dN/dS* models (Yang, 2006). By partitioning abstract substitution events into individual biophysical channels such as hydrophobicity, volume, and charge, we can define a more nuanced classification of molecular evolution. This resolution reveals that adaptation is often not a random exploration of sequence space, but rather a process of constrained diversification, where residues are driven to change in one property while being strictly prohibited from altering others. Conversely, PRIME identifies constraints at sites that appear evolutionarily neutral to rate-based models; at these positions, high turnover is permitted only among chemically similar residues, maintaining a functional constant despite sequence variation. In cases where the biophysical signal is more overt, such as the charge tuning observed in olfactory and immune receptors, the model explicitly identifies the directional shifts driving diversification.

Since its initial implementation as a part of the HyPhy package over 10 years ago, the PRIME framework—specifically S-PRIME—has been successfully applied across a diverse range of biological systems to resolve the specific physicochemical drivers of selection. These applications range from characterizing the adaptation of metabolic enzymes in extreme environments (Melo-Ferreira et al., 2014) and xenobiotic metabolism (Almeida et al., 2016) to dissecting the biophysical logic of viral drug resistance and immune escape (Grinev et al., 2016; Franzo et al., 2025; Shepherd et al., 2017). Furthermore, S-PRIME has identified critical constraints at protein-RNA interfaces (Mallik and Kundu, 2017) and distinguished between adaptive and neutral drivers of molecular evolution in both innate immune receptors (Velova et al., 2018) and plant populations (Lee-Yaw et al., 2019). Collectively, these studies underscore the utility of transforming abstract evolutionary rates into concrete mechanistic hypotheses across diverse taxonomic scales and functional paradigms.

This framework provides a connection between statistical phylogenetics and representation learning. While protein language models like ESM-2 (Lin et al., 2023) are robust predictors of fitness, the underlying drivers of their predictions remain difficult to interpret. The significant correlation between PRIME importance weights and the top principal components of PLM embeddings suggests that PRIME effectively characterizes the biophysical features learned by these models from substantial sequence databases. However, it is important to distinguish between interpretability and absolute predictive performance. While PLMs achieve higher zero-shot accuracy by capturing high-order sequence dependencies, PRIME provides a small-data alternative that resolves the mechanistic basis of selection from a single nucleotide sequence alignment. Moreover, by modeling physicochemical requirements rather than sequence frequency, PRIME reduces the impact of historical biases common to deep-learning approaches, identifying physicochemically viable substitutions that may be missing from training data but are permitted by the protein’s structural requirements.

Our simulation studies establish the statistical requirements for applying these biophysical models. The primary determinant of statistical power is the redundancy ratio (*R* = *N_subs_/N_aa_*), representing an approximation of how many times the evolutionary process has sampled the available state space. We find that *R >* 2.0 serves as a critical threshold for meaningful biophysical inference. Researchers must also account for the loss of signal under intense purifying selection, an information-theoretic blind spot where constraints paradoxically become difficult to detect by extinguishing the very mutational signal required for characterization. Consequently, the greatest potential for discovery lies at sites with moderate to high turnover but restricted chemical breadth.

However, despite its gains in biophysical realism, PRIME operates under several simplifying assumptions that define the practical boundaries of its application:

1. **Mutational Depth:** The site-level optimization of 5–10 parameters requires substantial evolutionary turnover. While the omnibus test serves as a necessary safeguard against overfitting, we find that the test is generally conservative for the bulk of sites with low informational redundancy (Figure 9C). Conversely, in regimes of extreme turnover with low diversity (toggling sites), the raw omnibus test can become anti-conservative, necessitating the methodical use of FDR correction (*q* ≤ 0.10) which effectively eliminates most of these artifacts (Figure 9B).
2. **Synonymous Rate Variation (SRV):** The current global (G-PRIME) and episodic (E-PRIME) implementations assume a constant synonymous rate (*α*) across the entire gene. For the genome-wide mammalian screen, we utilized G-PRIME with a global synonymous rate multiplier to maintain computational tractability, and direct comparabilty with traditional *dN/dS* methoods. While we acknowledge that unmodeled SRV can occasionally confound selection detection, our site-level S-PRIME implementation explicitly accounts for site-to-site variation, providing a more thorough validation for specific functional motifs where mutational hotspots may be a concern.
3. **Formulation of Positive Selection:** The functional form of the substitution rate (Equation 2) is mathematically optimized for purifying selection (*λ >* 0). In cases of diversifying selection (*λ <* 0), the rate increases with physicochemical distance. While we impose a computational ceiling on these rates to prevent numerical divergence, this formulation implies that adaptation favors maximizing chemical difference. Biologically, however, positive selection often seeks a new structural optimum rather than infinite divergence. Consequently, negative *λ* weights should be interpreted as a continuum ranging from the active selection for physicochemical change to the total relaxation of structural constraints. In regions like IDRs, a signature of ”diversification” may often reflect a permissive structural sieve that allows for neutral exploration of property space.
4. **Epistasis:** The current framework models each site as an independent entity. In reality, protein residues are highly cooperative, and the biophysical constraints at one site are contingent on the state of its neighbors. While S-PRIME effectively identifies the marginal selective pressures acting on individual residues, it does not explicitly model epistatic interactions. This limitation is shared by most site-specific frameworks, including Mutation-Selection (MutSel) models (Rodrigue et al., 2021), which characterize heterogeneous preferences across sites but typically assume site independence. Future work integrating PRIME’s biophysical grammar with hierarchical Bayesian models or explicit co-evolutionary frameworks could better resolve these structural dependencies and distinguish between alignment noise and genuine adaptation in rapidly evolving regions.
5. **Stationarity:** S-PRIME assumes a stationary process across the phylogeny. In contexts involving rapid selective sweeps or host-switching events, such as those observed in our H3N2 and HIV-RT analyses, this assumption represents a simplification of the underlying non-equilibrium dynamics. While the framework could be extended to allow for site-specific branch-to-branch variation via random-effects mixture modeling, such an implementation would necessitate a substantial increase in mutational depth to remain statistically identifiable. We have therefore opted for a practical trade-off that prioritizes site-level biophysical resolution, while offering E-PRIME as a complementary tool for detecting episodic fluctuations at the gene-wide level. Given that stationary models remain the standard for molecular evolutionary analysis and have consistently delivered biologically meaningful results, we view the current implementation as a robust first-order approximation of the physicochemical rules governing sequence transition.
6. **Property Collinearity:** Fundamental biophysical attributes are often correlated. Our analysis reveals high collinearity between hydrophobicity, volume, and polarity. While the omnibus test remains a robust detector of selection, the precise partitioning of selective signal between these dimensions remains subject to uncertainty. Researchers should interpret individual property weights as part of a synergistic biophysical profile rather than isolated effects.

Based on our performance benchmarks, we suggest a hierarchical workflow tailored to the specific research objective. G-PRIME can be used to obtain a rapid, gene-wide biophysical fingerprint of a protein family, while E-PRIME is suited for investigating lineage-specific adaptation or episodic property shifts. For resolving the mechanistic basis of specific functional or drug-resistance motifs at individual sites, S-PRIME should be applied to positions with sufficient mutational depth, defined by a redundancy ratio *R >* 2.0. For alignments with limited evolutionary information, simpler model configurations incorporating fewer properties are often worthwhile for resolving fundamental biophysical signals.

The ability to decode evolutionary history into concrete physical rules offers a powerful tool for prospective applications, such as identifying the biophysical constraints that channel viral immune escape or predicting the impact of mutations in uncharacterized proteins. By transforming the abstract study of evolutionary rates into a quantitative investigation of biophysical rules, PRIME provides a robust method for understanding the mechanistic drivers of genetic diversity.

## Acknowledgments and Disclaimers

This work was supported in part by the following awards: NIH/NHGRI (HG009299), NIH/NIGMS (GM151683), NSF (2419522), and NIH/NIAID (AI183870). llumina Inc. was not involved in and it does not sponsor or endorse this work; views expressed are those of the authors.

## 1 Computational Performance

We characterized the computational efficiency of the G-PRIME and E-PRIME implementations by analyzing the execution times for 18,944 mammalian genes. All analyses were performed on a standardized high-performance computing environment utilizing 4 cores of Ampere Altra (Neoverse-N1) ARM 128-core processors running at 3GHz. These benchmarks demonstrate that while the biophysical realism of PRIME requires more computational effort than property-agnostic models, the implementations remain practical for large-scale genomic applications. The G-PRIME model, which estimates gene-wide property weights, exhibits a mean execution time of 42.1 ± 82.2 minutes per gene. Over 75% of the mammalian alignments were processed in under 35 minutes, confirming that G-PRIME is suitable for routine gene-level characterization. These runtimes include LRT testing for each of the five individual property weights. The episodic E-PRIME model, which utilizes random-effects mixture modeling to capture fluctuating biophysical pressures, exhibits significantly higher computational demands. E-PRIME yielded a mean runtime of 4.57 ± 7.7 hours per gene, with a median of 2.36 hours. These runtimes also include LRT testing for each of the ten individual property weights (once per component). These results underscore that while S-PRIME and G-PRIME provide efficient site-level and gene-level resolution, the full characterization of episodic biophysical selection across the proteome necessitates substantial high-performance computing resources, totaling approximately 350,000 CPU-hours for the 120-mammal dataset.

### Implementation and Availability

The S-PRIME framework is implemented as a standard analysis within the HyPhy software package (version 2.5.94 or later). The G-PRIME and E-PRIME models are implemented using the FitModel analysis available in the hyphy-analyses repository (https://github.com/veg/hyphy-analyses).

Supporting scripts used for the analyses presented in this manuscript, including the mammalian genome screen, simulation pipelines, structural mapping, and DMS validation, are available in a dedicated reproducibility repository at https://github.com/veg/PRIME.

#### Running PRIME Models in HyPhy

PRIME models can be invoked via the HyPhy command-line interface. By default, the models utilize the 5 physico-chemical properties discussed in this manuscript: Hydrophobicity, Volume, Isoelectric Point, Alpha-Helix Propensity, and Beta-Sheet Propensity. Users may also provide custom property vectors or select from alternative sets (e.g., Atchley factors) during the analysis setup.

#### G-PRIME (Global Model)

Gene-wide physicochemical constraints are estimated using the FitModel.bf anal-ysis with the MG REV PROPERTIES model. This analysis is available in the hyphy-analyses repository:

hyphy [path_to_hyphy-analyses]/FitModel/FitModel.bf

--alignment [path_to_msa]

--model MG_REV_PROPERTIES

--property-set 5PROP

**E-PRIME (Episodic Model)** Episodic property selection is detected using the FitModel.bf analysis with the

MG REV PROPERTIES BSREL model:

hyphy [path_to_hyphy-analyses]/FitModel/FitModel.bf

--alignment [path_to_msa]

--model MG_REV_PROPERTIES_BSREL

--property-set 5PROP

**S-PRIME (Site-level Model)** Site-specific biophysical drivers are resolved using the dedicated prime analysis built into HyPhy:

hyphy prime --alignment [path_to_msa] --property-set 5PROP

#### Standard Output

All PRIME implementations generate a JSON-formatted result file containing maximum likelihood estimates for property weights (*λ*), model fit statistics (log-likelihood, AICc), and significance test results (LRT *p*-values). These files are designed for downstream visualization and integration into bioinformatics pipelines.

---

**Figure S1:**
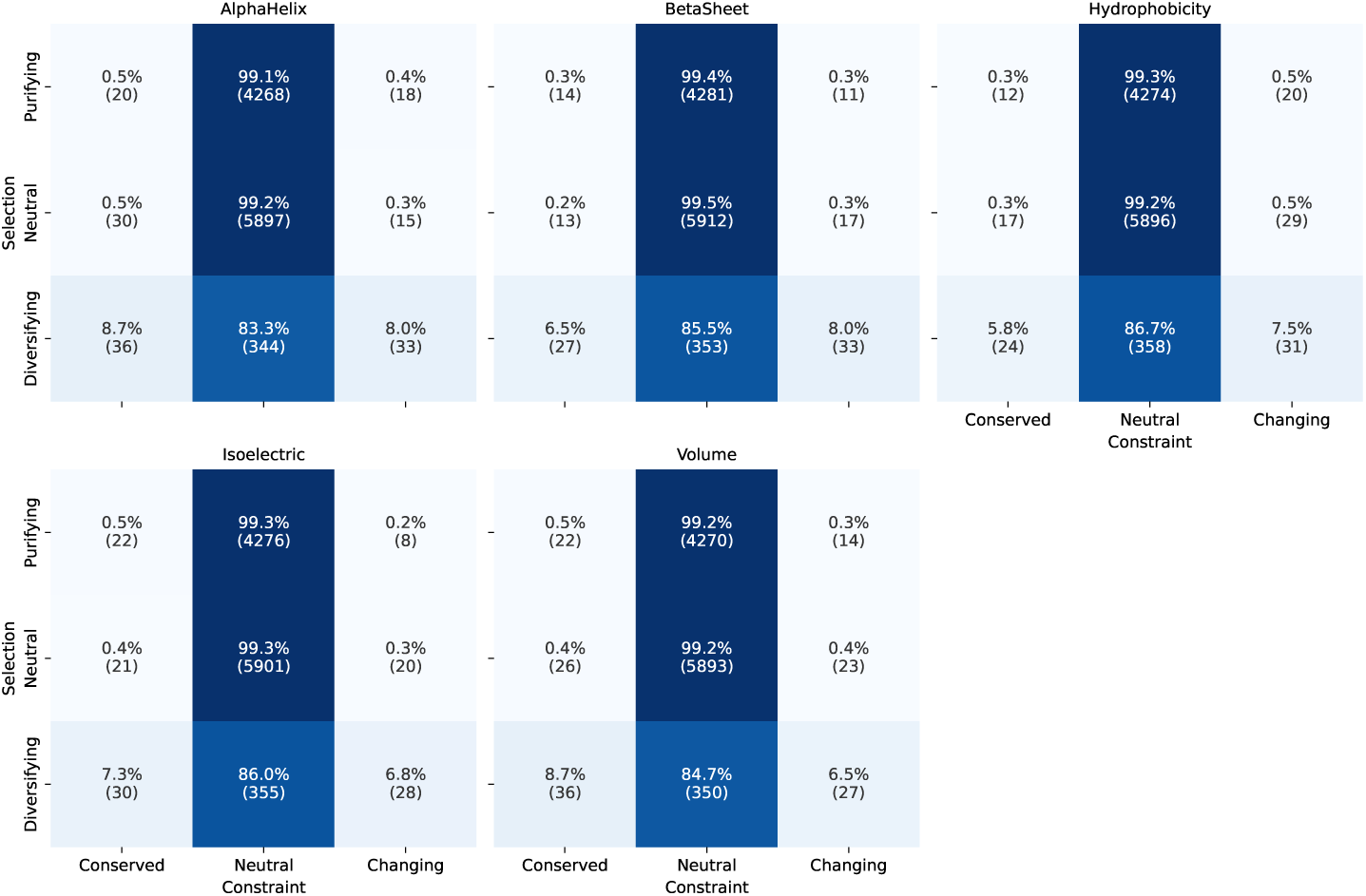
Conditional Probability of Physicochemical Constraints. Heatmap showing the percentage of sites within each selection class (Purifying, Neutral, Diversifying) that are Conserved, Neutral, or Changing for each physicochemical property. Raw site counts are shown in parentheses. Cryptic conservation is highlighted in the Neutral selection row, where 0.4% of sites exhibit significant property conservation despite lacking a rate-based signature of selection (*dN/dS* ≈ 1).

**Figure S2:**
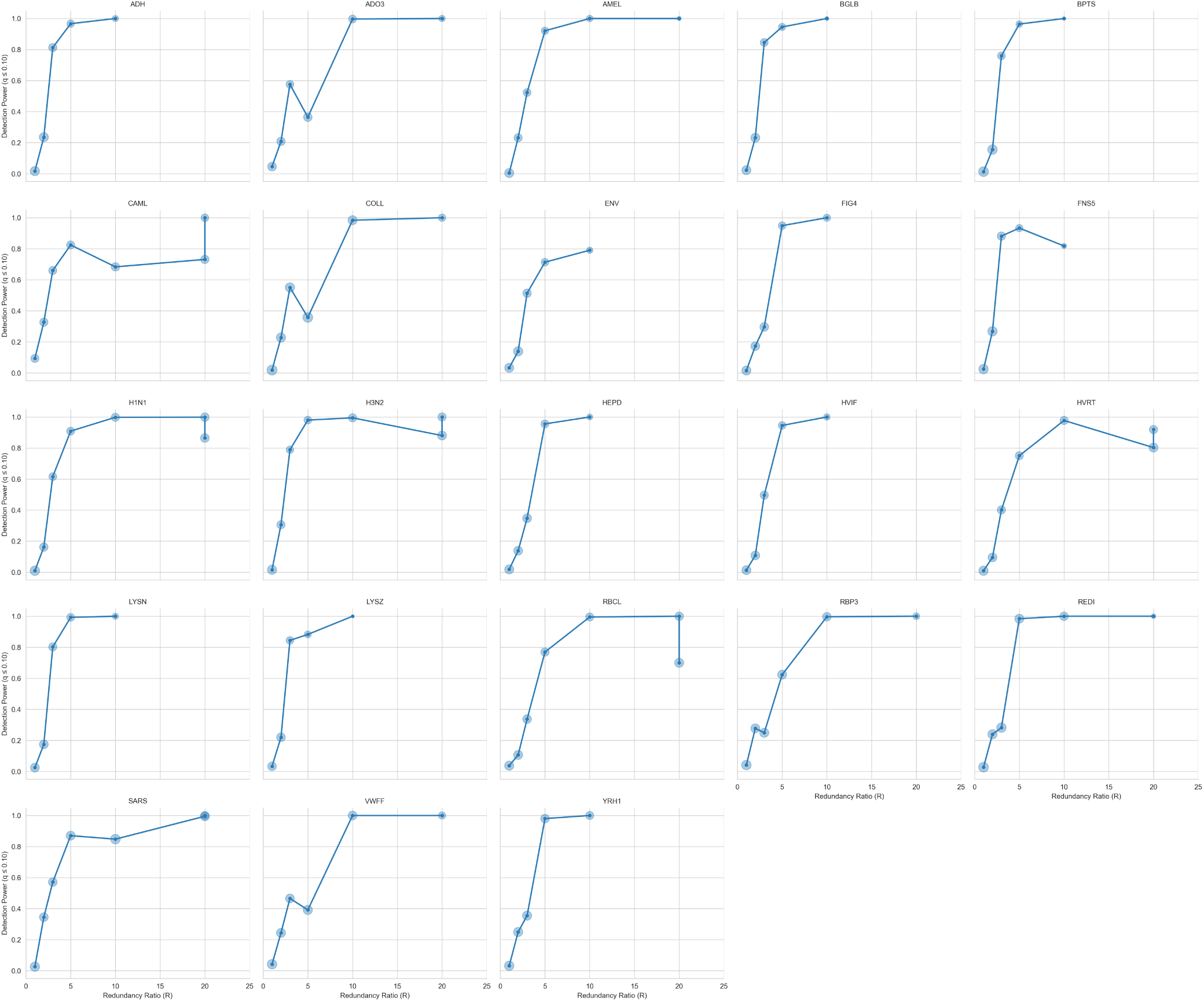
Detection power as a function of the redundancy ratio (*R*) for each of the 23 benchmark datasets. Detection is defined by the Benjamini-Hochberg corrected omnibus test (*q* ≤ 0.10). Bubbles denote the relative number of simulation sites in each *R* bin. The consistent non-linear scaling across all genes confirms that informational depth is the primary driver of sensitivity, regardless of taxonomic origin or structural context.

**Figure S3:**
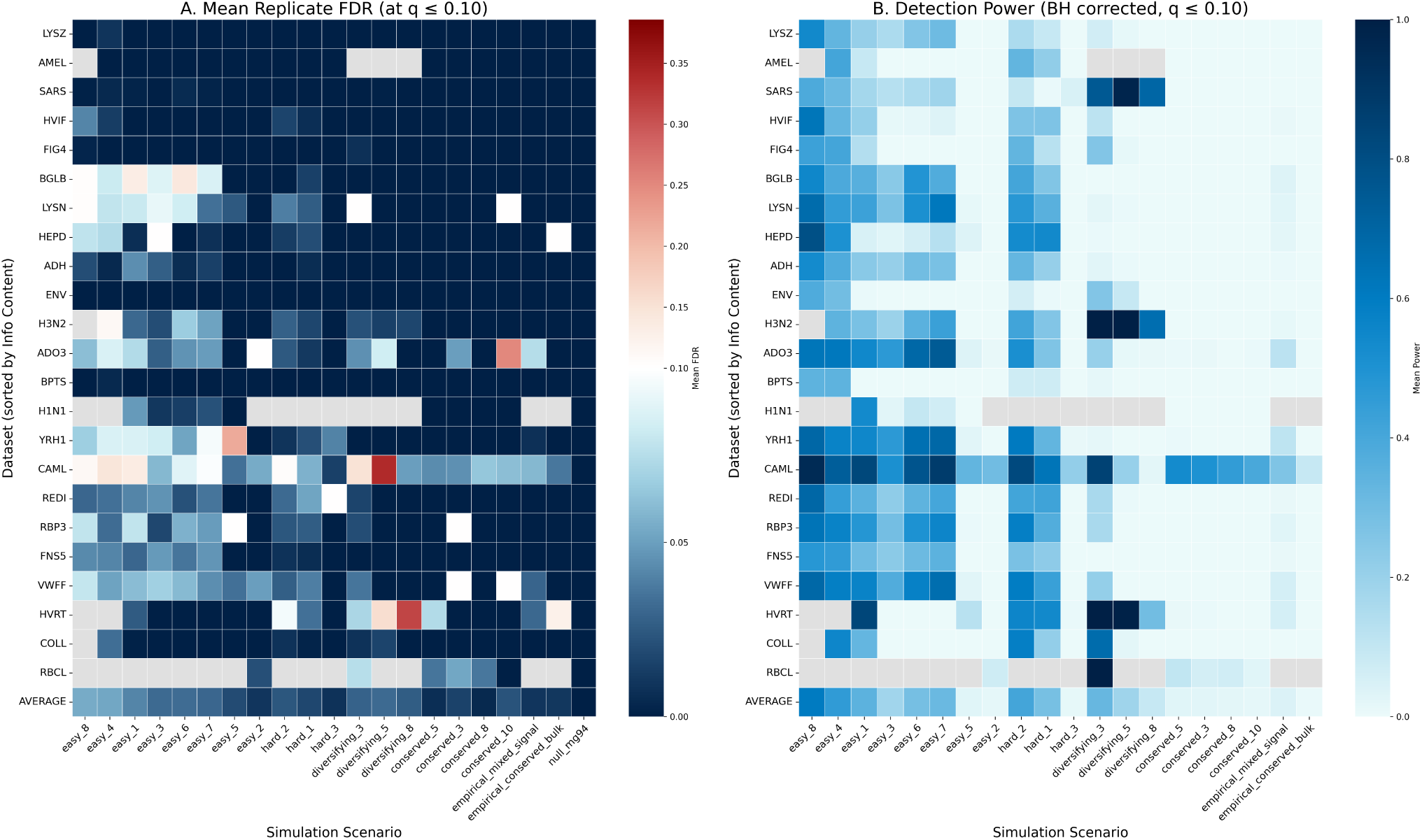
Dataset and scenario-specific performance S-PRIME performance on simulated data. (A) Empirical False Discovery Rate (FDR) calculated as the mean of replicate-level FDRs (*^F^ ^P^*) for each gene-scenario combination. The color scale is anchored at the nominal target *q* ≤ 0.10 (Vik palette). (B) Mean detection power after Benjamini-Hochberg correction. The Y-axis is sorted by informational content (*L* × *T*), ranging from minimal signal (LYSZ) to maximal signal (RBCL). Gray cells denote incomplete data (≤ 7 replicates). The final row (average) provides the scenario-wide mean across all benchmark genes.

**Table S1:**
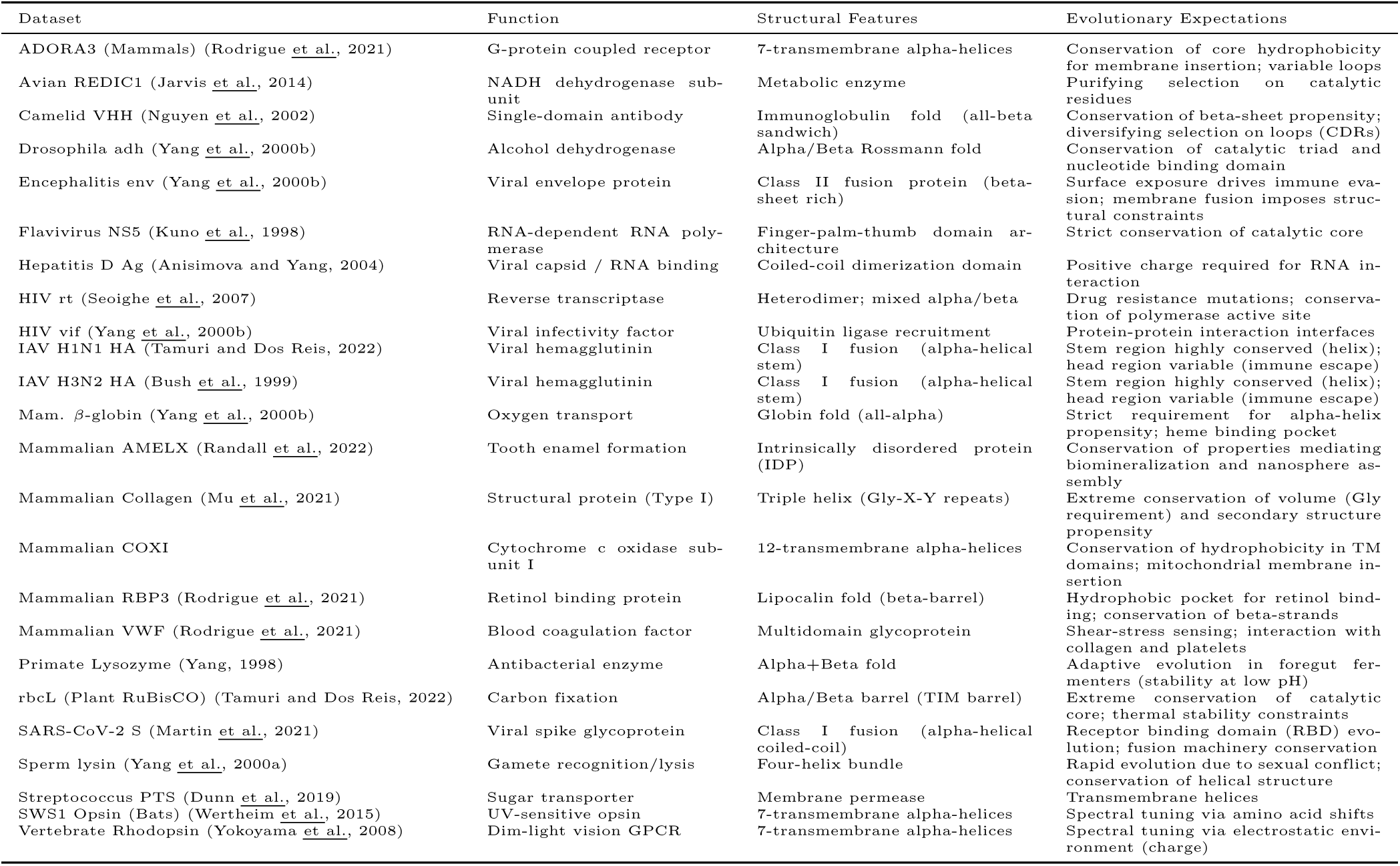
Functional and structural characteristics of the benchmark datasets.

**Table S2:**
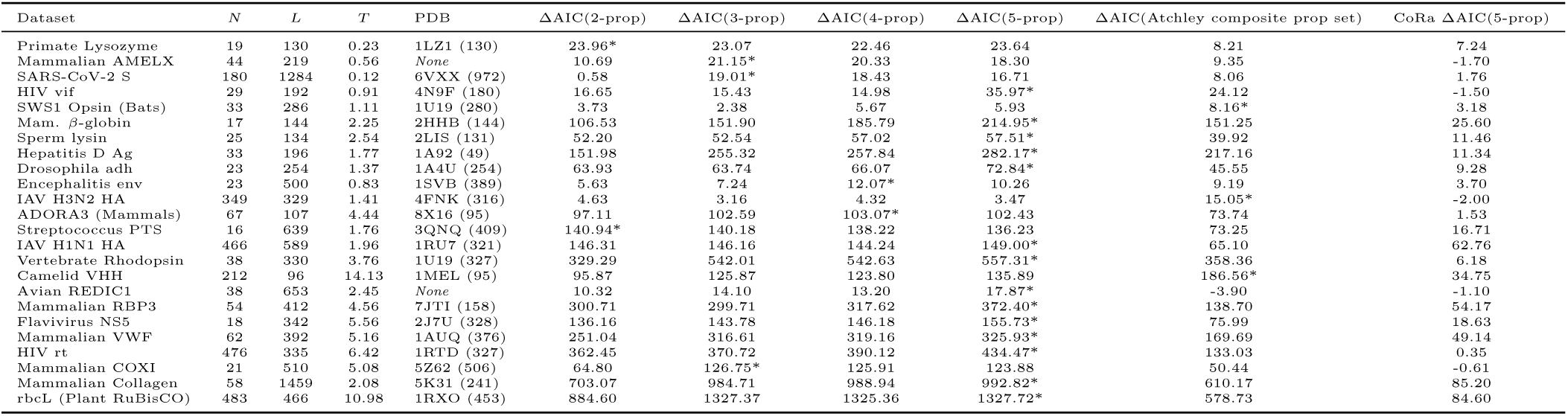
Model fit comparison and dataset characteristics for 24 benchmark alignments. For each dataset, we report the number of sequences (*N*), the number of codons (*L*), the total tree length (*T*, expected substitutions per site), and the PDB ID used for structural analysis (if available, with the number of mapped residues in parentheses). Datasets are sorted by the product of sequence length and tree length (*L* × *T*), a proxy for total evolutionary information. Fit improvement is shown as ΔAIC relative to the *MG*94×*REV* baseline (higher values indicate better fit). The best-fitting model for each dataset is marked with an asterisk (*) and highlighted in bold.

**Table S3:** Estimated property importance weights (*λ*) for the 5-prop PRIME model across 24 benchmark alignments. Datasets are sorted by the product of sequence length and tree length (*L* × *T*). *λ*s statistically significantly different from zero (Likelihood Ratio Test) are bolded and colored (Blue for conserved, Red for diversifying). Significance levels: * *p <* 0.05, ** *p <* 0.01, *** *p <* 0.001.

| Dataset | Hydrophobicity | Volume | Isoelectric Point | Alpha-Helix | Beta-Sheet |
| --- | --- | --- | --- | --- | --- |
| Primate Lysozyme | 0.55 | **0.94\*\*** | -0.22 | 0.19 | 0.59 |
| Mammalian AMELX | **0.13\*\*\*** | **0.35\*\*\*** | **0.37\*\*\*** | 0.12 | **0.01\*\*\*** |
| SARS-CoV-2 S | **0.19\*** | -0.05 | **-0.23\*\*\*** | 0.09 | 0.04 |
| HIV vif | 0.07 | **0.25\*\*** | -0.04 | 0.02 | 0.51 |
| SWS1 Opsin (Bats) | 0.04 | 0.12 | -0.02 | 0.15 | 0.15 |
| Mam. β-globin | **0.73\*\*\*** | **0.53\*\*\*** | **0.45\*\*\*** | **-0.48\*\*\*** | 0.66 |
| Sperm lysin | **0.21\*\*** | **0.39\*\*\*** | 0.07 | **-0.17\*** | 0.13 |
| Hepatitis D Ag | **0.35\*\*\*** | **0.43\*\*\*** | **0.45\*\*\*** | **-0.17\*\*** | 0.50 |
| Drosophila adh | **0.56\*\*\*** | 0.21 | 0.07 | **0.21\*** | 0.38 |
| Encephalitis env | **0.57\*\*** | 0.15 | **0.20\*** | **-0.32\*** | -0.07 |
| IAV H3N2 HA | 0.03 | **0.11\*** | -0.03 | 0.08 | 0.07 |
| ADORA3 (Mammals) | **0.51\*\*\*** | **0.26\*\*\*** | **0.12\*** | 0.09 | 0.10 |
| Streptococcus PTS | **0.43\*\*\*** | **0.72\*\*\*** | -0.06 | -0.02 | -0.01 |
| IAV H1N1 HA | **0.16\*\*\*** | **0.34\*\*\*** | -0.03 | -0.01 | 0.12 |
| Vertebrate Rhodopsin | **0.68\*\*\*** | **0.42\*\*\*** | **0.85\*\*\*** | -0.08 | 0.26 |
| Camelid VHH | **0.16\*\*\*** | **0.13\*\*\*** | **0.14\*\*\*** | 0.00 | 0.14 |
| Avian REDIC1 | **-0.12\*\*\*** | **0.08\*** | **0.05\*** | -0.04 | 0.09 |
| Mammalian RBP3 | **0.59\*\*\*** | **0.20\*\*\*** | -0.02 | **-0.20\*\*\*** | 0.32 |
| Flavivirus NS5 | **0.22\*\*** | **0.72\*\*\*** | **0.16\*\*** | **-0.16\*** | 0.28 |
| Mammalian VWF | **0.43\*\*\*** | **0.27\*\*\*** | **0.19\*\*\*** | **-0.09\*\*** | 0.11 |
| HIV rt | **0.20\*\*\*** | **0.72\*\*\*** | **0.06\*\*** | **-0.21\*\*\*** | 0.34 |
| Mammalian COXI | 0.10 | **0.83\*\*\*** | **0.74\*\*\*** | 0.09 | 0.01 |
| Mammalian Collagen | -0.01 | **1.20\*\*\*** | **0.53\*\*\*** | **-0.09\*\*** | **-0.10\*** |
| rbcL (Plant RuBisCO) | 0.02 | **0.58\*\*\*** | **0.35\*\*\*** | 0.01 | 0.05 |

**Table S4:** E-PRIME (5-prop) Parameter Estimates. For each dataset, we report the estimated property importance weights (*λ*) for the two mixture components. The weight of each component is shown in the second column. Values significantly different from zero (LRT *p <* 0.05) are shown in bold. Values at the upper bound of 10.00 are indicated with a dagger symbol.

| Dataset | Weight | Hydrophobicity | Volume | Isoelectric Pt | Alpha Helix | Beta Sheet |
| --- | --- | --- | --- | --- | --- | --- |
| Primate Lysozyme | 0.80 | -0.19 | -3.05 | -2.85 | 7.61 | 10.00<sup>†</sup> |
|  | 0.20 | 0.95 | 1.38 | 0.07 | -0.01 | 0.23 |
| SARS-CoV-2 S | 0.97 | 10.00<sup>†</sup> | -5.54 | -3.60 | 2.78 | 7.65 |
|  | 0.03 | 0.10 | 0.00 | -0.15 | 0.12 | 0.02 |
| HIV vif | 0.93 | 0.95 | 0.59 | 0.20 | -0.22 | 0.92 |
|  | 0.07 | -0.36 | -0.07 | -0.20 | 0.09 | 0.18 |
| SWS1 Opsin (Bats) | 0.79 | 2.18 | 8.31 | -0.18 | 10.00<sup>†</sup> | 1.47 |
|  | 0.21 | -0.03 | 0.04 | -0.03 | -0.06 | 0.05 |
| Mam. β-globin | 0.91 | 1.36 | 0.95 | 0.18 | 0.14 | 0.22 |
|  | 0.09 | 0.67 | 0.43 | 1.26 | -0.88 | 1.88 |
| Sperm lysin | 0.80 | 9.31 | -4.29 | 10.00<sup>†</sup> | 10.00<sup>†</sup> | 9.70 |
|  | 0.20 | 0.34 | 0.56 | 0.11 | -0.56 | 0.25 |
| Hepatitis D Ag | 0.90 | 3.48 | 4.25 | 2.83 | -1.92 | 1.02 |
|  | 0.10 | 0.31 | 0.21 | 0.55 | 0.22 | 0.50 |
| Drosophila adh | 0.94 | 10.00<sup>†</sup> | 1.69 | 10.00<sup>†</sup> | -0.10 | 0.76 |
|  | 0.06 | 0.62 | 0.34 | 0.05 | 0.21 | 0.64 |
| Encephalitis env | 0.62 | 5.11 | 10.00<sup>†</sup> | 5.58 | -1.15 | 2.02 |
|  | 0.38 | 0.61 | -0.02 | 0.15 | -0.30 | -0.13 |
| IAV H3N2 HA | 0.96 | 0.72 | 0.32 | -0.73 | -0.94 | 2.06 |
|  | 0.04 | -0.31 | 0.26 | 0.42 | 0.56 | -0.26 |
| ADORA3 (Mammals) | 0.71 | 9.00 | 0.74 | -0.04 | -3.48 | 6.88 |
|  | 0.29 | 0.35 | 0.40 | 0.11 | 0.42 | -0.12 |
| Streptococcus PTS | 0.87 | 10.00<sup>†</sup> | 0.40 | 0.61 | -0.20 | -0.41 |
|  | 0.13 | 0.60 | 1.37 | -0.12 | -0.14 | 0.11 |
| IAV H1N1 HA | 0.87 | -0.78 | 1.36 | 7.74 | 2.03 | -1.24 |
|  | 0.13 | 0.12 | 0.38 | -0.10 | -0.09 | 0.21 |
| Vertebrate Rhodopsin | 0.96 | 1.55 | 2.06 | 8.63 | -0.21 | 1.18 |
|  | 0.04 | 1.19 | 0.48 | 0.83 | -0.52 | 0.05 |
| Camelid VHH | 0.95 | 0.35 | 0.24 | 0.13 | -0.08 | 0.11 |
|  | 0.05 | -0.16 | -0.06 | 0.22 | 0.05 | 0.42 |
| Avian REDIC1 | 0.52 | -0.12 | 0.07 | 0.07 | -0.06 | 0.08 |
|  | 0.48 | 8.85 | 10.00<sup>†</sup> | -5.10 | 6.51 | 7.17 |
| Mammalian RBP3 | 0.82 | 2.04 | 0.64 | 0.83 | -1.56 | -0.75 |
|  | 0.18 | 0.37 | 0.03 | -0.28 | 0.57 | 0.89 |
| Flavivirus NS5 | 0.90 | -0.52 | 8.18 | 10.00<sup>†</sup> | 10.00<sup>†</sup> | 7.22 |
|  | 0.10 | 0.49 | 1.62 | 0.25 | -0.77 | 0.30 |
| Mammalian VWF | 0.73 | 8.89 | 1.50 | 10.00<sup>†</sup> | 0.29 | -0.60 |
|  | 0.27 | 0.40 | 0.33 | 0.16 | -0.15 | 0.12 |
| HIV rt | 0.96 | 10.00<sup>†</sup> | -0.54 | -0.55 | -4.84 | 2.34 |
|  | 0.04 | -0.03 | 1.12 | 0.17 | 0.00 | 0.21 |
| rbcL (Plant RuBisCO) | 0.97 | -1.74 | 7.90 | 10.00<sup>†</sup> | -0.70 | -0.33 |
|  | 0.03 | 0.28 | 0.29 | 0.31 | 0.03 | -0.04 |

**Table S5:** Selective Scenarios for S-PRIME Power Analysis. For each scenario, we report the biophysical importance factors (*λ*) applied to the 20% alternative partition. Properties: H: Hydrophobicity, V: Volume, P: Isoelectric Point (pI), *α*: Alpha-Helix, *β*: Beta-Sheet.

| ID | Description | $\lambda_H$ | $\lambda_V$ | $\lambda_P$ | $\lambda_\alpha$ | $\lambda_\beta$ |
| --- | --- | --- | --- | --- | --- | --- |
| easy_8 | Easy: hydro++ vol+ pol- helix- sheet- | 12.38 | 0.86 | -5.02 | -0.86 | -2.37 |
| easy_4 | Easy: hydro- vol++ pol++ helix- sheet++ | -4.91 | 3.85 | 1.16 | -2.37 | 12.42 |
| easy_1 | Easy: hydro- vol++ pol- helix++ sheet- | -1.05 | 2.20 | -0.66 | 11.37 | -3.50 |
| easy_3 | Easy: hydro++ vol- pol++ helix++ sheet- | 14.34 | -1.64 | 9.52 | 2.18 | -2.89 |
| easy_6 | Easy: hydro++ vol++ pol++ helix- sheet- | 2.21 | 13.22 | 2.96 | -4.16 | -0.35 |
| easy_7 | Easy: hydro++ vol- pol++ helix- sheet++ | 4.94 | -6.95 | 1.79 | -0.60 | 13.80 |
| easy_5 | Easy: hydro- vol++ pol++ helix+ sheet- | -1.61 | 1.98 | 13.23 | 0.83 | -0.87 |
| easy_2 | Easy: hydro+ pol+ helix- sheet+ | 0.95 | 0.09 | 0.63 | -0.38 | 0.25 |
| hard_2 | Hard: hydro+ vol+ pol+ helix- sheet+ | 0.55 | 1.19 | 5.73 | -2.56 | 1.07 |
| hard_1 | Hard: hydro+ vol+ pol- helix- sheet- | 2.99 | 3.44 | -1.69 | -0.58 | -0.99 |
| hard_3 | Hard: vol- pol+ helix+ sheet+ | 0.15 | -0.55 | 0.54 | 1.00 | 0.40 |
| diversifying_3 | Diversifying: lambda=-3.0 | 0.00 | -3.00 | 0.00 | 0.00 | 0.00 |
| diversifying_5 | Diversifying: lambda=-5.0 | 0.00 | -5.00 | 0.00 | 0.00 | 0.00 |
| diversifying_8 | Diversifying: lambda=-8.0 | 0.00 | -8.00 | 0.00 | 0.00 | 0.00 |
| conserved_5 | Conservation: lambda=5.0 | 0.00 | 5.00 | 0.00 | 0.00 | 0.00 |
| conserved_3 | Conservation: lambda=3.0 | 0.00 | 3.00 | 0.00 | 0.00 | 0.00 |
| conserved_8 | Conservation: lambda=8.0 | 0.00 | 8.00 | 0.00 | 0.00 | 0.00 |
| conserved_10 | Conservation: lambda=10.0 | 0.00 | 10.00 | 0.00 | 0.00 | 0.00 |
| empirical_mixed_signal | Empirical Cluster 4: Strong Sheet conservation, moderate Vol diversification | -1.26 | -1.99 | 3.41 | 0.27 | 9.39 |
| empirical_conserved_bulk | Empirical Cluster 1: Moderate conservation on multiple props | 2.99 | 2.93 | 2.25 | -0.14 | -0.96 |
| null_mg94 | Strict Null: All lambdas = 0 (MG94) | 0.00 | 0.00 | 0.00 | 0.00 | 0.00 |

**Table S6:**
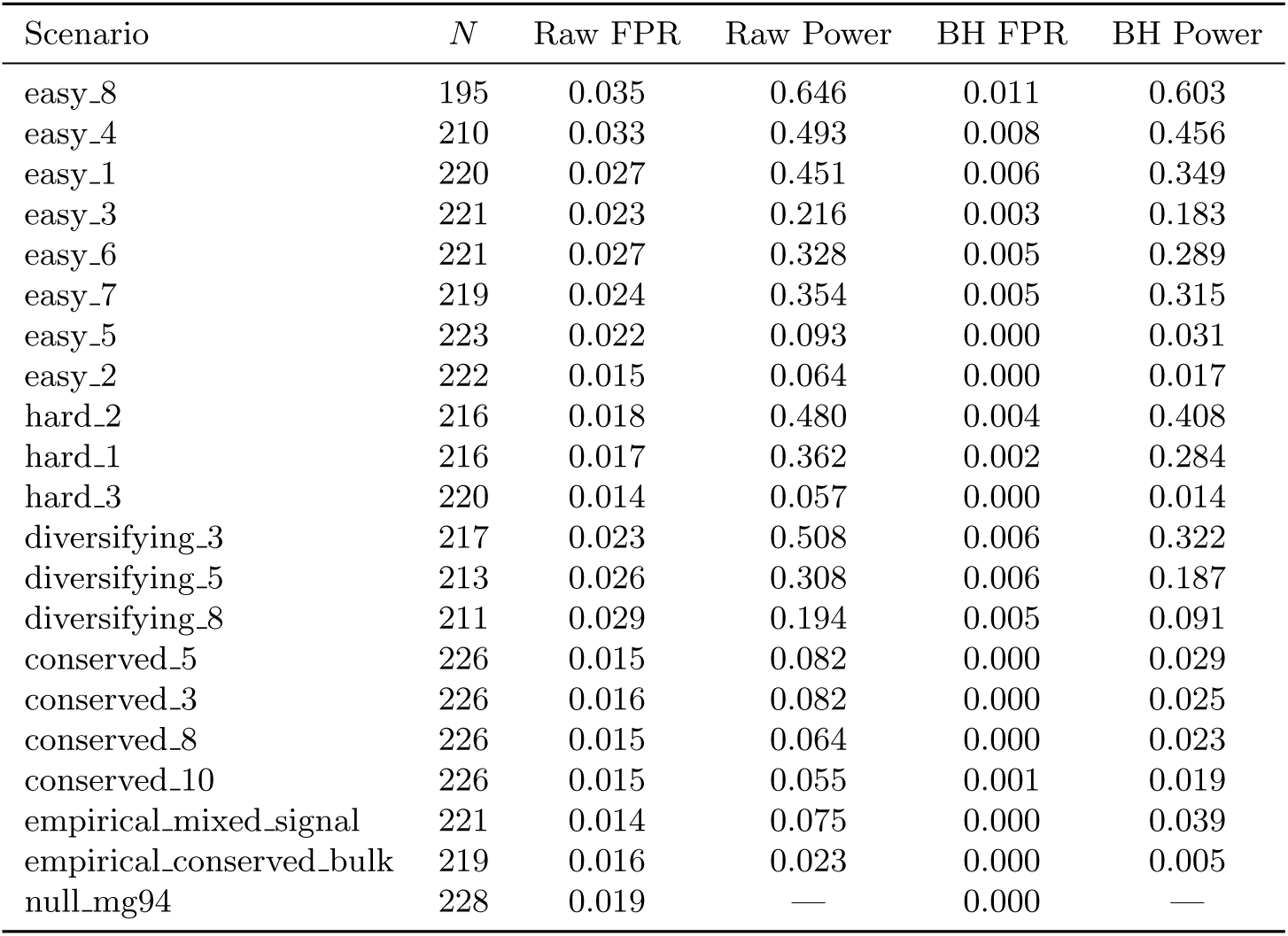
S-PRIME Simulation Results by selective scenario. We report the mean False Positive Rate (FPR) and Power (sensitivity) averaged over all replicates (*N*) for both raw (*α* = 0.05) and Benjamini-Hochberg corrected (*q* ≤ 0.10) omnibus tests. For the null mg94 scenario, where all sites are neutral, results from both partitions are pooled into a single FPR estimate.

**Table S7:** Detailed S-PRIME Performance and Descriptive Statistics by Dataset for Scenario easy_1 (*λ* ≈ 11). *R* = *N_subs_/N_aa_* denotes the Redundancy Ratio. FPR and Power are reported for raw (*α* = 0.05) and BH-corrected (*q* ≤ 0.10) tests.

| Dataset | $N$ | FPR (Raw) | FPR (BH) | Power (Raw) | Power (BH) | Med $R_{null}$ | Med $R_{alt}$ | Med $S_{null}$ | Med $S_{alt}$ |
| --- | --- | --- | --- | --- | --- | --- | --- | --- | --- |
| AMELX | 10 | 0.002 | 0.000 | 0.286 | 0.091 | 0.5 | 1.4 | 1 | 8 |
| Bacterial_PTS_trehalose_transporter_suIII | 10 | 0.002 | 0.000 | 0.107 | 0.002 | 0.5 | 0.8 | 1 | 4 |
| COL1A1 | 10 | 0.005 | 0.000 | 0.478 | 0.329 | 0.5 | 1.7 | 1 | 9 |
| ENCenv | 10 | 0.000 | 0.000 | 0.126 | 0.005 | 0.0 | 1.0 | 0 | 3 |
| Fig4E | 10 | 0.001 | 0.000 | 0.302 | 0.138 | 0.5 | 1.3 | 1 | 6 |
| HIV_RT | 10 | 0.022 | 0.006 | 0.885 | 0.834 | 1.3 | 5.1 | 6 | 48 |
| HIVvif | 10 | 0.007 | 0.000 | 0.395 | 0.213 | 0.7 | 1.5 | 2 | 8 |
| HepatitisD | 10 | 0.018 | 0.001 | 0.287 | 0.048 | 0.8 | 1.9 | 3 | 19 |
| IAV-human-H1N1-HA | 8 | 0.031 | 0.006 | 0.604 | 0.538 | 1.0 | 2.5 | 3 | 16 |
| InfluenzaA | 8 | 0.032 | 0.003 | 0.335 | 0.280 | 1.0 | 0.5 | 3 | 1 |
| REDIC1 | 10 | 0.029 | 0.004 | 0.438 | 0.352 | 1.0 | 1.0 | 4 | 2 |
| SARS-CoV-2-spike | 9 | 0.003 | 0.000 | 0.177 | 0.172 | 0.0 | 0.0 | 0 | 0 |
| adh | 10 | 0.037 | 0.003 | 0.325 | 0.239 | 0.8 | 0.5 | 3 | 1 |
| adora3 | 10 | 0.046 | 0.012 | 0.605 | 0.545 | 1.7 | 1.3 | 11 | 4 |
| bglobin | 10 | 0.070 | 0.015 | 0.514 | 0.362 | 1.1 | 1.2 | 7 | 3 |
| camelid | 10 | 0.072 | 0.037 | 0.830 | 0.825 | 3.7 | 6.6 | 41 | 26 |
| flavNS5 | 10 | 0.036 | 0.003 | 0.428 | 0.307 | 1.0 | 1.0 | 4 | 2 |
| lysin | 10 | 0.043 | 0.009 | 0.559 | 0.430 | 1.2 | 1.2 | 7 | 4 |
| lysozyme | 10 | 0.006 | 0.000 | 0.265 | 0.208 | 0.5 | 0.5 | 1 | 1 |
| rbcL | 5 | 0.050 | 0.015 | 0.726 | 0.698 | 3.0 | 3.1 | 24 | 12 |
| rbp3 | 10 | 0.040 | 0.010 | 0.559 | 0.482 | 1.4 | 1.3 | 8 | 4 |
| vwf | 10 | 0.042 | 0.010 | 0.618 | 0.567 | 1.6 | 1.5 | 10 | 5 |
| yokoyama.rh1.cds.mod.1-990 | 10 | 0.046 | 0.013 | 0.639 | 0.548 | 1.4 | 1.5 | 10 | 5 |

**Table S8:** Site Classification Counts across Benchmark Datasets. Sites were classified based on the decision tree in Figure 6, integrating MEME selection detection (*p* ≤ 0.05) with PRIME physicochemical structure (FDR ≤ 0.10). *Adaptive* classes (left) correspond to sites under diversifying selection; *Neutral/Purifying* classes (right) correspond to sites without rate elevation. Abbreviations: Constr. (Constrained), Div. (Diversification), Unconstr. (Unconstrained), Drv. (Drivers), Neut. (Neutral).

| Dataset | Adaptive (MEME +) |  |  |  | Neutral / Purifying (MEME -) |  |  |  |
| --- | --- | --- | --- | --- | --- | --- | --- | --- |
|  | Constr. Div. | Driven Div. | Complex Div. | Unconstr. Div. | Cryptic Constr. | Cryptic Drv. | Complex Neut. | Non-Sig. |
| rbcL (Plant RuBisCO) | 47 | 10 | 15 | 11 | 43 | 11 | 41 | 288 |
| Mammalian Collagen | 9 | 3 | 7 | 30 | 15 | 5 | 15 | 1375 |
| Mammalian VWF | 4 | 0 | 2 | 1 | 30 | 4 | 28 | 323 |
| HIV rt | 12 | 3 | 4 | 3 | 14 | 7 | 19 | 273 |
| Vertebrate Rhodopsin | 1 | 2 | 0 | 9 | 15 | 8 | 24 | 271 |
| Camelid VHH | 8 | 4 | 8 | 9 | 14 | 2 | 11 | 40 |
| Avian REDIC1 | 0 | 0 | 1 | 45 | 2 | 1 | 1 | 603 |
| Sperm lysin | 10 | 0 | 3 | 13 | 9 | 3 | 7 | 89 |
| Hepatitis D Ag | 3 | 0 | 4 | 12 | 5 | 1 | 5 | 166 |
| IAV H1N1 HA | 4 | 1 | 2 | 17 | 1 | 1 | 1 | 562 |
| IAV H3N2 HA | 2 | 0 | 3 | 12 | 2 | 1 | 4 | 305 |
| Mammalian RBP3 | 0 | 0 | 1 | 3 | 2 | 3 | 8 | 395 |
| Flavivirus NS5 | 1 | 0 | 0 | 1 | 7 | 1 | 3 | 329 |
| HIV vif | 0 | 0 | 0 | 8 | 1 | 1 | 3 | 179 |
| ADORA3 (Mammals) | 0 | 0 | 0 | 1 | 4 | 2 | 3 | 97 |
| Mam. beta-globin | 0 | 0 | 0 | 0 | 8 | 1 | 1 | 134 |
| SARS-CoV-2 S | 0 | 0 | 0 | 7 | 2 | 0 | 0 | 1275 |
| Streptococcus PTS | 0 | 0 | 0 | 9 | 0 | 0 | 0 | 630 |
| Drosophila adh | 0 | 0 | 0 | 8 | 0 | 0 | 0 | 246 |
| Mammalian COXI | 0 | 0 | 0 | 2 | 0 | 1 | 0 | 507 |
| SWS1 Opsin (Bats) | 0 | 0 | 0 | 3 | 0 | 0 | 0 | 283 |
| Encephalitis env | 0 | 0 | 0 | 1 | 1 | 0 | 0 | 498 |
| Mammalian AMELX | 0 | 0 | 0 | 2 | 0 | 0 | 0 | 217 |
| Primate Lysozyme | 0 | 0 | 0 | 0 | 0 | 0 | 0 | 130 |

**Table S9:**
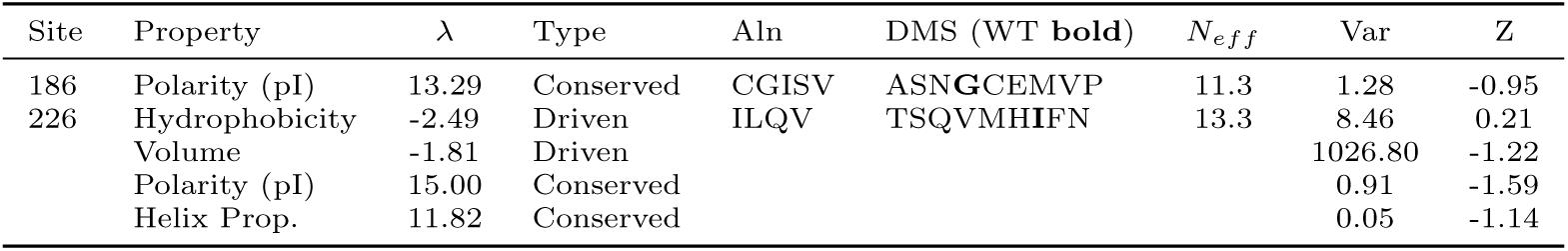
PRIME vs. DMS Validation for Significant H3N2 Sites. Properties shown are individually significant after the full two-stage hierarchical correction (*q <* 0.05, controlling for selective inference across the entire gene). ’DMS Residues’ account for 80 percent of the preference mass. A negative Z-score confirms clustering in property space (Constraint). Consistent with Figure 7, sites located in the buried core (e.g., Site 226) exhibit higher importance weights and more restricted experimental variance.

## References

1. Almeida, D., Maldonado, E., Khan, I., Silva, L., Gilbert, M. T. P., Zhang, G., Jarvis, E. D., O’Brien, S. J., Johnson, W. E., and Antunes, A. 2016. Whole-genome identification, phylogeny, and evolution of the cytochrome p450 family 2 (cyp2) subfamilies in birds. Genome Biol Evol, 8(4): 1115–31.

2. Anisimova, M. and Yang, Z. 2004. Molecular evolution of the hepatitis delta virus antigen gene: recombination or positive selection? J Mol Evol, 59(6): 815–26.

3. Atchley, W. R., Zhao, J., Fernandes, A. D., and Druke, T. 2005. Solving the protein sequence metric problem. Proceedings of the National Academy of Sciences of the United States of America, 102(18): 6395–6400.

4. Benjamini, Y. and Bogomolov, M. 2014. Selective inference on multiple families of hypotheses. Journal of the Royal Statistical Society Series B: Statistical Methodology, 76(1): 297–318.

5. Bloom, J. D., Drummond, D. A., Arnold, F. H., and Wilke, C. O. 2006. Structural determinants of the rate of protein evolution in yeast. Molecular biology and evolution, 23(8): 1751–1761.

6. Bloom, J. D., Raval, A., and Wilke, C. O. 2007. Thermodynamics of neutral protein evolution. Genetics, 175(1): 255–266.

7. Bush, R. M., Bender, C. A., Subbarao, K., Cox, N. J., and Fitch, W. M. 1999. Predicting the evolution of human influenza a. Science, 286(5446): 1921–5.

8. Chou, P. Y. and Fasman, G. D. 1974. Prediction of protein conformation. Biochemistry, 13(2): 222–245.

9. Conant, G. C., Wagner, G. P., and Stadler, P. F. 2007. Modeling amino acid substitution patterns in orthologous and paralogous genes. Mol Phylogenet Evol, 42(2): 298–307.

10. Delport, W., Scheffler, K., and Seoighe, C. 2008. Frequent toggling between alternative amino acids is driven by selection in hiv-1. PLoS Pathog, 4(12): e1000242.

11. Dill, K. A. 1990. Dominant forces in protein folding. Biochemistry, 29(31): 7133–7155.

12. Dunker, A. K., Lawson, J. D., Brown, C. J., Williams, R. M., Romero, P., Oh, J. S., Oldfield, C. J., Campen, A. M., Ratliff, C. M., Hipps, K. W., et al. 2001. Intrinsically disordered protein. Journal of molecular graphics and modelling, 19(1): 26–59.

13. Dunn, K. A., Kenney, T., Gu, H., and Bielawski, J. P. 2019. Improved inference of site-specific positive selection under a generalized parametric codon model when there are multinucleotide mutations and multiple nonsynonymous rates. BMC Evol Biol, 19(1): 22.

14. Durbin, R., Eddy, S., Krogh, A., and Mitchison, G. 1998. Biological Sequence Analysis: Probabilistic Models of Proteins and Nucleic Acids. Cambridge University Press.

15. Felsenstein, J. 2004. Inferring Phylogenies. Sinauer Associates.

16. Franzo, G., Legnardi, M., Poletto, F., Baston, R., Cecchinato, M., Drigo, M., and Tucciarone, C. M. 2025. Host-driven evolution of pcv2: insights into genetic diversity and adaptation. Front Immunol, 16: 1577436.

17. Goldman, N. and Yang, Z. 1994. A codon-based model of nucleotide substitution for protein-coding DNA sequences. Mol. Biol. Evol., 11: 725–736.

18. Grantham, R. 1974. Amino acid difference formula to help explain protein evolution. Science, 185(4154): 862–864.

19. Grinev, A., Chancey, C., Volkova, E., Anez, G., Heisey, D. A. R., Winkelman, V., Foster, G. A., Williamson, P., Stramer, S. L., and Rios, M. 2016. Genetic variability of west nile virus in u.s. blood donors from the 2012 epidemic season. PLoS Negl Trop Dis, 10(5): e0004717.

20. Hecker, N. and Hiller, M. 2019. A genome alignment of 120 mammals highlights ultraconserved element stability and placental-specific enhancers. Nature Communications, 10(1): 4811.

21. Hurvich, C. M. and Tsai, C.-L. 1989. Regression and time series model selection in small samples. Biometrika, 76(2): 297–307.

22. Jarvis, E. D., Mirarab, S., Aberer, A. J., Li, B., Houde, P., Li, C., Ho, S. Y. W., Faircloth, B. C., Nabholz, B., Howard, J. T., Suh, A., Weber, C. C., da Fonseca, R. R., Li, J., Zhang, F., Li, H., Zhou, L., Narula, N., Liu, L., Ganapathy, G., Boussau, B., Bayzid, M. S., Zavidovych, V., Subramanian, S., Gabaldon, T., Capella-Gutierrez, S., Huerta-Cepas, J., Rekepalli, B., Munch, K., Schierup, M., Lindow, B., Warren, W. C., Ray, D., Green, R. E., Bruford, M. W., Zhan, X., Dixon, A., Li, S., Li, N., Huang, Y., Derryberry, E. P., Bertelsen, M. F., Sheldon, F. H., Brumfield, R. T., Mello, C. V., Lovell, P. V., Wirthlin, M., Schneider, M. P. C., Prosdocimi, F., Samaniego, J. A., Vargas Velazquez, A. M., Alfaro-Nunez, A., Campos, P. F., Petersen, B., Sicheritz-Ponten, T., Pas, A., Bailey, T., Scofield, P., Bunce, M., Lambert, D. M., Zhou, Q., Perelman, P., Driskell, A. C., Shapiro, B., Xiong, Z., Zeng, Y., Liu, S., Li, Z., Liu, B., Wu, K., Xiao, J., Yinqi, X., Zheng, Q., Zhang, Y., Yang, H., Wang, J., Smeds, L., Rheindt, F. E., Braun, M., Fjeldsa, J., Orlando, L., Barker, F. K., Jønsson, K. A., Johnson, W., Koepfli, K.-P., O’Brien, S., Haussler, D., Ryder, O. A., Rahbek, C., Willerslev, E., Graves, G. R., Glenn, T. C., McCormack, J., Burt, D., Ellegren, H., Alstrom, P., Edwards, S. V., Stamatakis, A., Mindell, D. P., Cracraft, J., Braun, E. L., Warnow, T., Jun, W., Gilbert, M. T. P., and Zhang, G. 2014. Whole-genome analyses resolve early branches in the tree of life of modern birds. Science, 346(6215): 1320–31.

23. Jones, C. T., Youssef, N., Susko, E., and Bielawski, J. P. 2020. A phenotype–genotype codon model for detecting adaptive evolution. Systematic biology, 69(4): 722–738.

24. Jumper, J., Evans, R., Pritzel, A., Green, T., Figurnov, M., Ronneberger, O., Tunyasuvunakool, K., Bates, R., Zıdek, A., Potapenko, A., Bridgland, A., Meyer, C., Kohl, S. A. A., Ballard, A. J., Cowie, A., Romera-Paredes, B., Nikolov, S., Jain, R., Adler, J., Back, T., Petersen, S., Reiman, D., Clancy, E., Zielinski, M., Steinegger, M., Pacholska, M., Berghammer, T., Bodenstein, S., Silver, D., Vinyals, O., Senior, A. W., Kavukcuoglu, K., Kohli, P., and Hassabis, D. 2021. Highly accurate protein structure prediction with alphafold. Nature, 596(7873): 583–589.

25. Kabsch, W. and Sander, C. 1983. Dictionary of protein secondary structure: pattern recognition of hydrogen-bonded and geometrical features. Biopolymers: Original Research on Biomolecules, 22(12): 2577–2637.

26. Kosakovsky Pond, S. and Frost, S. D. 2005. Not so different after all: A comparison of methods for detecting amino acid sites under selection. Mol. Biol. Evol., 22: 1208–1222.

27. Kosakovsky Pond, S. L., Poon, A. F. Y., Brown, A. J. L., Frost, S. D. W., and Muse, S. V. 2008. A maximum likelihood method for detecting directional evolution in protein sequences and its application to influenza A virus. Mol. Biol. Evol., 25: 1809–1824.

28. Kuno, G., Chang, G. J., Tsuchiya, K. R., Karabatsos, N., and Cropp, C. B. 1998. Phylogeny of the genus flavivirus. J Virol, 72(1): 73–83.

29. Kyte, J. and Doolittle, R. F. 1982. A simple method for displaying the hydropathic character of a protein. Journal of Molecular Biology, 157(1): 105–132.

30. Lacerda, M., Scheffler, K., and Seoighe, C. 2010. Epitope discovery with phylogenetic hidden markov models. Molecular Biology and Evolution, 27(5): 1212–1220.

31. Lee, J. M., Huddleston, J., Doud, M. B., Hooper, K. A., Wu, N. C., Bedford, T., and Bloom, J. D. 2018. Deep mutational scanning of hemagglutinin helps predict evolutionary fates of influenza virus variants. Proceedings of the National Academy of Sciences, 115(35): E8276–E8285.

32. Lee-Yaw, J. A., Grassa, C. J., Joly, S., Andrew, R. L., and Rieseberg, L. H. 2019. An evaluation of alternative explanations for widespread cytonuclear discordance in annual sunflowers (helianthus). New Phytol, 221(1): 515–526.

33. Liberles, D. A., Teichmann, S. A., Bahar, I., Bastolla, U., Bloom, J., Bornberg-Bauer, E., Colwell, L. J., de Koning, J., Dokholyan, I., Echave, J., et al. 2012. The interface of protein structure, protein biophysics, and molecular evolution. Protein Science, 21(6): 769–785.

34. Lin, Z., Akin, H., Rao, R., Hie, B., Zhu, Z., Lu, W., Smetanin, D., Verkuil, R., Kabeli, O., Shmueli, Y., et al. 2023. Evolutionary-scale prediction of atomic-level protein structure with a language model. Science, 379(6637): 1123–1130.

35. Lucaci, A. G., Zehr, J. D., Enard, D., Thornton, J. W., and Kosakovsky Pond, S. L. 2023. Evolutionary shortcuts via multinucleotide substitutions and their impact on natural selection analyses. Mol Biol Evol, 40(7): msad150.

36. Mallik, S. and Kundu, S. 2017. Modular organization of residue-level contacts shapes the selection pressure on individual amino acid sites of ribosomal proteins. Genome Biol Evol, 9(4): 916–931.

37. Martin, D. P., Weaver, S., Tegally, H., San, J. E., Shank, S. D., Wilkinson, E., Lucaci, A. G., Giandhari, J., Naidoo, S., Pillay, Y., Singh, L., Lessells, R. J., NGS-SA, COVID-19 Genomics UK (COG-UK), Gupta, R. K., Wertheim, J. O., Nekturenko, A., Murrell, B., Harkins, G. W., Lemey, P., MacLean, O. A., Robertson, D. L., de Oliveira, T., and Kosakovsky Pond, S. L. 2021. The emergence and ongoing convergent evolution of the sars-cov-2 n501y lineages. Cell, 184(20): 5189–5200.e7.

38. Melo-Ferreira, J., Vilela, J., Fonseca, M. M., da Fonseca, R. R., Boursot, P., and Alves, P. C. 2014. The elusive nature of adaptive mitochondrial dna evolution of an arctic lineage prone to frequent introgression. Genome Biol Evol, 6(4): 886–96.

39. Mirny, L. A. and Shakhnovich, E. I. 1999. Universally conserved positions in protein folds: reading evolutionary signals about stability, folding kinetics and function. Journal of molecular biology, 291(1): 177–199.

40. Miyata, T., Miyazawa, S., and Yasunaga, T. 1979. Two types of amino acid substitutions in protein evolution. Journal of molecular evolution, 12(3): 219–236.

41. Mu, Y., Tian, R., Xiao, L., Sun, D., Zhang, Z., Xu, S., and Yang, G. 2021. Molecular evolution of tooth-related genes provides new insights into dietary adaptations of mammals. Journal of Molecular Evolution, 89(7): 458–471.

42. Murrell, B., Wertheim, J. O., Moola, S., Weighill, T., Scheffler, K., and Kosakovsky Pond, S. L. 2012a. Detecting individual sites subject to episodic diversifying selection. PLoS Genet, 8(7): e1002764.

43. Murrell, B., Wertheim, J. O., Moodley, S., Murrell, K., and Pond, S. L. K. 2012b. Detecting individual sites subject to episodic diversifying selection. PLoS genetics, 8(7): e1002764.

44. Murrell, B., de Oliveira, T., Seebregts, C., Kosakovsky Pond, S. L., Scheffler, K., and on behalf of the Southern African Treatment and Resistance Network (SATuRN) Consortium 2012c. Modeling hiv-1 drug resistance as episodic directional selection. PLoS Comput Biol, 8(5): e1002507.

45. Murrell, B., Weaver, S., Smith, M. D., Wertheim, J. O., Murrell, S., Aylward, A., Eren, K., Pollner, T., Martin, D. P., Smith, D. M., Scheffler, K., and Pond, S. L. K. 2015. Gene-wide identification of episodic selection. Molecular Biology and Evolution, 32(5): 1365–1371.

46. Muse, S. V. and Gaut, B. S. 1994. A likelihood approach for comparing synonymous and nonsynonymous nucleotide substitution rates, with application to the chloroplast genome. Mol. Biol. Evol., 11: 715–724.

47. Ngandu, N. K., Seoighe, C., and Scheffler, K. 2009. Evidence of hiv-1 adaptation to host hla alleles following chimp-to-human transmission. Virol J, 6: 164.

48. Nguyen, V. K., Su, C., Muyldermans, S., and van der Loo, W. 2002. Heavy-chain antibodies in camelidae; a case of evolutionary innovation. Immunogenetics, 54(1): 39–47.

49. Randall, J. G., Gatesy, J., and Springer, M. S. 2022. Molecular evolutionary analyses of tooth genes support sequential loss of enamel and teeth in baleen whales (mysticeti). Molecular Phylogenetics and Evolution, 171: 107463.

50. Richards, F. M. 1974. The interpretation of protein structures: total volume, group volume distributions and packing density. Journal of molecular biology, 82(1): 1–14.

51. Rives, A., Meier, J., Sercu, T., Goyal, S., Lin, Z., Liu, J., Guo, D., Ott, M., Zitnick, C. L., Ma, J., et al. 2021. Biological structure and function emerge from scaling unsupervised learning to 250 million protein sequences. Proceedings of the National Academy of Sciences, 118(15): e2016239118.

52. Rodrigue, N., Latrille, T., and Lartillot, N. 2021. A bayesian mutation-selection framework for detecting site-specific adaptive evolution in protein-coding genes. Mol Biol Evol, 38(3): 1199–1208.

53. Rogers, G. and Paulson, J. 1983. Receptor determinants of human and animal influenza virus isolates: differences in receptor specificity of the h3 hemagglutinin based on sialic acid linkage. Virology, 127(2): 361–373.

54. Sainudiin, R., Wong, W. S. W., Yogeeswaran, K., Nasrallah, J. B., Yang, Z., and Nielsen, R. 2005. Detecting site-specific physicochemical selective pressures: applications to the class i hla of the human major histocompatibility complex and the srk of the plant sporophytic self-incompatibility system. J Mol Evol, 60(3): 315–26.

55. Seoighe, C., Ketwaroo, F., Pillay, V., Scheffler, K., Wood, N., Duffet, R., Zvelebil, M., Martinson, N., McIntyre, J., Morris, L., and Hide, W. 2007. A model of directional selection applied to the evolution of drug resistance in hiv-1. Molecular Biology and Evolution, 24(4): 1025–1031.

56. Serohijos, A. W. and Shakhnovich, E. I. 2014. Merging molecular evolution and biophysics: from molecules to organisms. Current opinion in structural biology, 26: 84–91.

57. Shepherd, F. K., Murtaugh, M. P., Chen, F., Culhane, M. R., and Marthaler, D. G. 2017. Longitudinal surveillance of porcine rotavirus b strains from the united states and canada and in silico identification of antigenically important sites. Pathogens, 6(4): 64.

58. Shrake, A. and Rupley, J. 1973. Environment and exposure to solvent of protein atoms. lysozyme and insulin. Journal of molecular biology, 79(2): 351–371.

59. Sugiura, N. 1978. Further analysts of the data by akaike’ s information criterion and the finite corrections. Communications in Statistics-theory and Methods, 7(1): 13–26.

60. Tamuri, A. U. and Dos Reis, M. 2022. A mutation-selection model of protein evolution under persistent positive selection. Mol Biol Evol, 39(1): msab309.

61. Tien, M. Z., Meyer, A. G., Sydykova, D. K., Spielman, S. J., and Wilke, C. O. 2013. Maximum allowed solvent accessibilites of residues in proteins. PloS one, 8(11): e80635.

62. Tokuriki, N. and Tawfik, D. S. 2009. Stability effects of mutations and protein evolvability. Current opinion in structural biology, 19(5): 596–604.

63. Velova, H., Gutowska-Ding, M. W., Burt, D. W., and Vinkler, M. 2018. Toll-like receptor evolution in birds: Gene duplication, pseudogenization, and diversifying selection. Mol Biol Evol, 35(9): 2170–2184.

64. Weber, C. C. and Whelan, S. 2019. Physicochemical amino acid properties better describe substitution rates in large populations. Mol Biol Evol, 36(4): 679–690.

65. Wertheim, J. O., Murrell, B., Smith, M. D., Kosakovsky Pond, S. L., and Scheffler, K. 2015. Relax: detecting relaxed selection in a phylogenetic framework. Mol Biol Evol, 32(3): 820–32.

66. Wong, W. S. W., Sainudiin, R., and Nielsen, R. 2006. Identification of physicochemical selective pressure on protein encoding nucleotide sequences. BMC Bioinformatics, 7: 148.

67. Xia, X. and Li, W.-H. 1998. What amino acid properties affect protein evolution? Journal of molecular evolution, 47(5): 557–564.

68. Yang, Z. 1998. Likelihood ratio tests for detecting positive selection and application to primate lysozyme evolution. Mol Biol Evol, 15(5): 568–73.

69. Yang, Z. 2006. Computational Molecular Evolution. Oxford University Press.

70. Yang, Z., Swanson, W. J., and Vacquier, V. D. 2000a. Maximum-likelihood analysis of molecular adaptation in abalone sperm lysin reveals variable selective pressures among lineages and sites. Mol Biol Evol, 17(10): 1446–55.

71. Yang, Z. H., Nielsen, R., Goldman, N., and Pedersen, A. M. K. 2000b. Codon-substitution models for heterogeneous selection pressure at amino acid sites. Genetics, 155: 431–449.

72. Yokoyama, S., Tada, T., Zhang, H., and Britt, L. 2008. Elucidation of phenotypic adaptations: Molecular analyses of dim-light vision proteins in vertebrates. Proc Natl Acad Sci U S A, 105(36): 13480–5.

73. Zamyatnin, A. A. 1972. Protein volume in solution. Progress in Biophysics and Molecular Biology, 24: 107–123.

74. Zarin, T., Tsai, C. N., Ba, A. N., and Moses, A. M. 2019. Selection maintains a specific molecular signature in the intrinsically disordered regions of proteins. eLife, 8: e48175.

